# Cross-disorder analysis of schizophrenia and 19 immune diseases reveals genetic correlation

**DOI:** 10.1101/068684

**Authors:** Jennie G Pouget, Schizophrenia Working Group of the Psychiatric Genomics Consortium, Buhm Han, Yang Wu, Emmanuel Mignot, Hanna M Ollila, Jonathan Barker, Sarah Spain, Nick Dand, Richard Trembath, Javier Martin, Maureen D Mayes, Lara Bossini-Castillo, Elena López-Isac, Ying Jin, Stephanie A Santorico, Richard A Spritz, Soumya Raychaudhuri, Jo Knight

**Affiliations:** Campbell Family Mental Health Research Institute, Centre for Addiction and Mental Health, Toronto, ON, Canada; Institute of Medical Sciences, University of Toronto, Toronto, ON, Canada; Department of Psychiatry, University of Toronto, Toronto, ON, Canada; Department of Biomedical Sciences, Seoul National University College of Medicine, Seoul, Republic of Korea; Institute for Molecular Bioscience, The University of Queensland, Brisbane, QLD 4072, Australia; Center for Sleep Sciences and Medicine, Department of Psychiatry and Behavioral Sciences, Stanford University, School of Medicine, Palo Alto, CA, USA; Finnish Institute for Molecular Medicine, Helsinki, Finland; Center for Genomic Medicine, Massachusetts General Hospital, Boston, Massachusetts, USA and Broad Institute, Cambridge, Massachusetts, USA; School of Basic and Medical Biosciences, Faculty of Life Sciences and Medicine, King’s College London, UK; St. John’s Institute of Dermatology, Faculty of Life Sciences and Medicine, King’s College London, London, UK; Wellcome Trust Sanger Institute, Wellcome Trust Genome Campus, Hinxton, Cambridge, UK; Queen Mary University of London, Barts and the London School of Medicine and Dentistry, London, UK; Institute of Parasitology and Biomedicine López-Neyra, IPBLN-CSIC, Granada, Spain; The University of Texas Health Science Center–Houston, Houston, TX, USA; Human Medical Genetics and Genomics Program, University of Colorado School of Medicine, Aurora, CO, USA; Department of Pediatrics, University of Colorado School of Medicine, Aurora, CO, USA; Department of Mathematical and Statistical Sciences, University of Colorado Denver, Denver, CO, USA; Department of Biostatistics and Informatics, Colorado School of Public Health, University of Colorado, Aurora, CO, USA; Department of Medicine, Brigham and Women's Hospital and Harvard Medical School, Boston, MA, USA; Partners HealthCare Center for Personalized Genetic Medicine, Boston, MA, USA; Program in Medical and Population Genetics, Broad Institute of MIT and Harvard, Cambridge, MA, USA; Division of Genetics, Brigham and Women’s Hospital, Harvard Medical School, Boston, MA, USA; Division of Rheumatology, Brigham and Women’s Hospital, Harvard Medical School, Boston, MA, USA; Faculty of Medical and Human Sciences, University of Manchester, Manchester, UK; Lancaster Medical School and Data Science Institute, Lancaster University, Lancaster, UK

## Abstract

Epidemiological studies indicate that many immune diseases occur at different rates among people with schizophrenia compared to the general population. Here, we evaluated whether this phenotypic correlation between immune diseases and schizophrenia might be explained by shared genetic risk factors (**genetic correlation**). We used data from a large genome-wide association study (GWAS) of schizophrenia (N=35,476 cases and 46,839 controls) to compare the genetic architecture of schizophrenia to 19 immune diseases. First, we evaluated the association with schizophrenia of 581 variants previously reported to be associated with immune diseases at genome-wide significance. We identified three variants with pleiotropic effects, located in regions associated with both schizophrenia and immune disease. Our analyses provided the strongest evidence of pleiotropy at rs1734907 (∼85kb upstream of *EPHB4*), a variant which was associated with increased risk of both Crohn’s disease (OR = 1.16, P = 1.67×10^−13^) and schizophrenia (OR = 1.07, P = 7.55×10^−6^). Next, we investigated genome-wide sharing of common variants between schizophrenia and immune diseases using polygenic risk scores (PRS) and cross-trait LD Score regression (LDSC). PRS revealed significant genetic overlap with schizophrenia for narcolepsy (p=4.1×10^−4^), primary biliary cirrhosis (p=1.4×10^−8^), psoriasis (p=3.6×10^−5^), systemic lupus erythematosus (p=2.2×10^−8^), and ulcerative colitis (p=4.3×10^−4^). Genetic correlations between these immune diseases and schizophrenia, estimated using LDSC, ranged from 0.10 to 0.18 and were consistent with the expected phenotypic correlation based on epidemiological data. We also observed suggestive evidence of sex-dependent genetic correlation between schizophrenia and multiple sclerosis (interaction p=0.02), with genetic risk scores for multiple sclerosis associated with greater risk of schizophrenia among males but not females. Our findings suggest that shared genetic risk factors contribute to the epidemiological co-occurrence of schizophrenia and certain immune diseases, and suggest that in some cases this genetic correlation is sex-dependent.

**Author Summary:** Immune diseases occur at different rates among patients with schizophrenia compared to the general population. While the reasons for this phenotypic correlation are unclear, shared genetic risk (**genetic correlation**) has been proposed as a contributing factor. Prior studies have estimated the genetic correlation between schizophrenia and a handful of immune diseases, with conflicting results. Here, we performed a comprehensive cross-disorder investigation of schizophrenia and 19 immune diseases. We identified three individual genetic variants associated with both schizophrenia and immune diseases, including a variant near *EPHB4* – a gene whose protein product guides the migration of lymphocytes towards infected cells in the immune system and the migration of neuronal axons in the brain. We demonstrated significant genome-wide genetic correlation between schizophrenia and narcolepsy, primary biliary cirrhosis, psoriasis, systemic lupus erythematosus, and ulcerative colitis. Finally, we identified a potential sex-dependent pleiotropic effect between schizophrenia and multiple sclerosis. Our findings point to shared genetic risk for schizophrenia and at least a subset of immune diseases, which likely contributes to their epidemiological co-occurrence. These results raise the possibility that the same genetic variants may exert their effects on neurons or immune cells to influence the development of psychiatric and immune disorders, respectively.

## Introduction

Despite recent advances in identifying key biomarkers and genetic loci for schizophrenia, its pathophysiology remains poorly understood [1, 2]. One interesting epidemiological observation is that the risk of developing many immune-mediated diseases is increased among patients with schizophrenia [3–5], and vice versa [6, 7]. Here, we use the term **immune disease** to broadly encompass both autoimmune and inflammatory disorders. While there are discrepancies among studies regarding which immune diseases are most strongly correlated with schizophrenia, there is converging evidence that these diseases co-occur at a greater rate than is expected by chance [3–7]. A notable exception is rheumatoid arthritis (RA), where a consistent inverse association with schizophrenia has been observed [5, 8].

Genetic factors have long been proposed as an explanation for the differing prevalence of immune diseases among patients with schizophrenia compared to the general population [5, 6]. The recently reported role of *complement component 4* (*C4*) variation in schizophrenia [9] illustrates a potential shared genetic mechanism in the development of immune and psychiatric disorders. Genetic variants conferring increased *C4* expression protect against developing systemic lupus erythematosus (SLE), possibly by increased tagging of apoptotic cells – which are the trigger for autoantibody development in SLE – leading to more effective clearance by macrophages [10]. The same genetic mechanism may increase the risk of developing schizophrenia, by increased tagging of neuronal synapses for elimination by microglia leading to excessive synaptic pruning [9]. We hypothesize that similar shared genetic mechanisms may occur throughout the genome, with cellular manifestations in immune cells and neurons influencing the development of immune and psychiatric disorders, respectively. Previously, we found that susceptibility to schizophrenia does not appear to be driven by the broad set of loci harboring immune genes [11]. However, not all genetic variants conferring risk of immune disease fall within immune loci. Here, we evaluated whether common genetic variants influencing the risk of 19 different immune diseases may also be involved in schizophrenia.

Our cross-disorder genetic approach is supported by recent successes in identifying shared genetic risk variants (**pleiotropy**) across a variety of human diseases [12–18]. Pleiotropy is emerging as a pervasive phenomenon in the human genome [19–21], and cross-disorder studies characterizing the nature of genotype-phenotype relationships have the potential to yield significant insights into disease etiology. For instance, cross-trait genetic analyses have shed new light on cardiovascular disease and lipid biology – and shifted attention away from HDL as a potential treatment target – by demonstrating that increased HDL cholesterol levels do not reduce the risk of myocardial infarction [14]. In psychiatry, cross-disorder analyses have identified significant pleiotropy between schizophrenia, bipolar disorder, and major depressive disorder, indicating that these diseases are not as distinct at a pathophysiological level as current diagnostic criteria suggest [12, 13, 22].

While previous studies have investigated genome-wide pleiotropy between schizophrenia and immune disorders, results have been inconsistent (**S1 Table**). Genetic correlation has been reported between schizophrenia and Crohn’s disease [23–27], multiple sclerosis [28], primary biliary cirrhosis [25], psoriasis [29], rheumatoid arthritis [23, 24], systemic lupus erythematosus [24, 25], and type 1 diabetes [23, 24, 26, 27] in some studies, but not in others [8, 13, 16, 24, 30]. Interestingly, negative genetic correlation (whereby genetic risk protects against developing schizophrenia) has also been reported for RA [31], in keeping with the inverse epidemiological association [5, 8].

Additional studies are needed to reconcile the inconsistencies in existing cross-trait analyses of schizophrenia and immune disorders, with careful attention towards potential confounding variables (e.g. population stratification, linkage disequilibrium, non-independence of genome-wide association study (GWAS) samples, and sex-specific effects). To this end we have performed a comprehensive cross-disorder analysis of schizophrenia and 19 immune diseases, using data from the largest available genetic studies. Our findings add to a growing body of literature supporting pervasive pleiotropy between schizophrenia and immune diseases. We extend existing literature by including 10 immune diseases that have not previously been compared with schizophrenia, prioritizing pleiotropic genes through integrative analyses of multiomics data, estimating how much of the phenotypic correlation between schizophrenia and immune diseases was explained by the genetic correlations we observed, and providing novel evidence for potential sex-specific pleiotropy between schizophrenia and immune disease.

## Results

### Defining immune risk variants

We identified immune-mediated diseases with robust GWAS findings using ImmunoBase (http://www.immunobase.org; accessed 7 June 2015), an online resource providing curated GWAS data for immune-related human diseases. These included the following 19 diseases: alopecia areata (AA), ankylosing spondylitis (AS), autoimmune thyroid disease (ATD), celiac disease (CEL), Crohn’s disease (CRO), inflammatory bowel disease (IBD), juvenile idiopathic arthritis (JIA), multiple sclerosis (MS), narcolepsy (NAR), primary biliary cirrhosis (PBC), primary sclerosing cholangitis (PSC), psoriasis (PSO), rheumatoid arthritis (RA), Sjögren’s syndrome (SJO), systemic lupus erythematosus (SLE), systemic sclerosis (SSC), type 1 diabetes (T1D), ulcerative colitis (UC), and vitiligo (VIT). Notably, the majority of IBD risk variants were also risk variants for CRO and/or UC. For 14 of these immune diseases (see **Table 1**), we also obtained full GWAS or Immunochip summary statistics allowing us to conduct additional polygenic risk scoring (PRS) [30, 32] and cross-trait Linkage Disequilibrium Score regression (LDSC) analyses [16].

**Table 1.** Description of datasets analyzed

|  |  |  |  |  |  | Total number of SNPs |  |  |
| --- | --- | --- | --- | --- | --- | --- | --- | --- |
|  | Abr | Genome-wide significant SNPs<sup>a</sup> | Polygenic risk scoring<sup>b</sup> | Cases | Controls | Full GWAS | Merged with SCZ<sup>c</sup> | Pruned<sup>d</sup> |
| Schizophrenia | SCZ | - | Target [1] | 35,476 | 46,839 | - | - | - |
| Height | HGT | - | Negative control [43] | 253,288 | - | 2,085,602 | 2,035,446 | 124,888 |
| Alopecia areata | AA | 11 | - | - | - | - | - | - |
| Ankylosing spondylitis | AS | 23 | - | - | - | - | - | - |
| Autoimmune thyroid disease | ATD | 7 | - | - | - | - | - | - |
| Celiac disease | CEL | 38 | Training [68] | 12,041 | 12,228 | 133,352<sup>f</sup> | 90,922 | 19,698 |
| Crohn's disease | CRO | 119 | Training [69] | 5,956 | 14,927 | 12,276,506 | 4,990,991 | 114,950 |
| Inflammatory bowel disease | IBD | 145 | Training [69] | 12,882 | 21,770 | 12,716,150 | 5,095,448 | 116,346 |
| Juvenile idiopathic arthritis | JIA | 22 | Training [70] | 772<sup>e</sup> | 8,530<sup>e</sup> | 122,330<sup>f</sup> | 98,477 | 20,337 |
| Multiple sclerosis | MS | 103 | Training [36] | 14,498 | 24,091 | 155,756<sup>f</sup> | 108,118 | 21,818 |
| Narcolepsy | NAR | 3 | Training [71] | 1,886 | 10,421 | 109,768<sup>f</sup> | 92,859 | 19,866 |
| Primary biliary cirrhosis | PBC | 19 | Training [75] | 2,764 | 10,475 | 1,038,537 | 1,041,977 | 97,806 |
| Primary sclerosing cholangitis | PSC | 12 | - | - | - | - | - | - |
| Psoriasis | PSO | 34 | Training [76] | 2,178 | 5,175 | 7,586,779 | 3,701,354 | 107,002 |
| Rheumatoid arthritis | RA | 77 | Training [72] | 5,539 | 20,169 | 2,090,825 | 2,087,383 | 126,049 |
| Sjögren's syndrome | SJO | 6 | - | - | - | - | - | - |
| Systemic lupus erythematosus | SLE | 19 | Training [73] | 4,036 | 6,959 | 7,915,251 | 6,539,217 | 264,374 |
| Systemic sclerosis | SSC | 4 | Training [77] | 1,486<sup>g</sup> | 3,477<sup>g</sup> | 253,179<sup>f</sup> | 251,441 | 66,402 |
| Type 1 diabetes | T1D | 56 | Training [74] | 9,340<sup>h</sup> | 12,835 | 123,081<sup>f</sup> | 98,418 | 20,835 |
| Ulcerative colitis | UC | 96 | Training [69] | 6,968 | 20,464 | 12,255,263 | 5,167,266 | 120,720 |
| Vitiligo | VIT | 16 | Training [78] | 1,381 | 14,518 | 8,790,155 | 6,223,502 | 257,654 |

<sup>a</sup>We obtained lists of genome-wide significant SNPs for each autoimmune disease from ImmunoBase, and processed them as described in **Supplementary Methods**; <sup>b</sup>The following columns provide details for datasets used in the polygenic risk scoring analysis. We used effect sizes

Given that human leukocyte antigen (HLA) alleles within the major histocompatibility complex (MHC) region (chromosome 6: 25-34 Mb) account for a significant proportion of heritability of immune and inflammatory disorders [33], we considered HLA and non-HLA risk variants separately in our analyses. Within the MHC region we considered only the most strongly associated HLA variant (including SNPs, imputed HLA amino acid sites, and classical alleles) for each disease based on univariate analysis in previously published studies (see **Table 2**), because multivariate conditional analyses reporting adjusted effect sizes of independent HLA variants were not available for all immune diseases. Outside of the MHC region, we considered all non-HLA variants curated in ImmunoBase for each of the 19 immune diseases.

**Table 2.** Association of top HLA variants for immune diseases in schizophrenia

| Disease | HLA variant | Autoimmune | | Schizophrenia | | $r^2$ with top SCZ SNP<sup>a</sup> |
| --- | --- | --- | --- | --- | --- | --- |
| | | $p$ | OR | $p$ | OR | |
| **AA [92]** | **HLA-DRB1#37Asn** | **$4.99 \times 10^{-73}$** | **0.42** | **$4.85 \times 10^{-9}$** | **0.91** | **0.04** |
| AS [93] | HLA-B*27 | $<1 \times 10^{-100}$ | 46 | 0.13 | 1.05 | 0 |
| ATD [94] | rs2281388<br>(tags HLA-DPB1*05:01) | $1.50 \times 10^{-65}$ | 1.64 | 0.39 | 1.04<sup>b</sup> | 0 |
| **CEL [95]** | **HLA-DQB1#74Ala** | **n.r.** | **2.14** | **$2.16 \times 10^{-12}$** | **0.89** | **0.11** |
| CRO [96] | HLA-DRB1*01:03 | $3.00 \times 10^{-62}$ | 2.51 | 0.61 | 0.96 | 0 |
| IBD [96] | HLA-DRB1*01:03 | $1.93 \times 10^{-112}$ | 3.01 | 0.61 | 0.96 | 0 |
| JIA [70] | rs7775055 | $3.14 \times 10^{-174}$ | 6.01 | 0.12 | 0.94 | 0 |
| MS [97] | HLA-DRB1*15:01 | $1.40 \times 10^{-234}$ | 2.92 | $5.10 \times 10^{-3}$ | 1.06 | 0 |
| NAR [98] | HLA-DQB1*06:02 | $1.04 \times 10^{-120}$ | 251 | $7.30 \times 10^{-3}$ | 1.06 | 0 |
| PBC [99] | HLA-DQA1*04:01 | $5.90 \times 10^{-45}$ | 3.06 | 0.20 | 0.95 | 0 |
| **PSC [100]** | **HLA-B\*08:01** | **$3.70 \times 10^{-246}$** | **2.82** | **$5.65 \times 10^{-16}$** | **0.84** | **0.2** |
| PSO [101] | HLA-C*06:02 | $2.10 \times 10^{-201}$ | 3.26 | 0.55 | 0.99 | 0 |
| RA [102] | HLA-DRB1#11Val | $<1 \times 10^{-581}$ | 3.80 | $2.68 \times 10^{-4}$ | 1.07 | 0 |
| **SJO [103]** | **HLA-DQB1\*02:01** | **$1.38 \times 10^{-95}$** | **3.36** | **$3.84 \times 10^{-15}$** | **0.85** | **0.11** |
| SLE [104] | HLA-DRB1#13Arg | $7.99 \times 10^{-10}$ | 1.55<sup>c</sup> | $5.81 \times 10^{-4}$ | 1.07 | 0 |
| SSC [105] | rs17500468 (TAP2) | $5.87 \times 10^{-62}$ | 2.87 | $6.76 \times 10^{-4}$ | 1.07 | 0 |
| T1D [106] | HLA-DQB1#57Ala | $<1 \times 10^{-1000}$ | 5.17 | $7.80 \times 10^{-4}$ | 0.95 | 0.06 |
| UC [96] | rs6927022 | $8.00 \times 10^{-154}$ | 1.49 | $3.37 \times 10^{-4}$ | 1.06 | 0.03 |
| VIT [78] | rs9271597<br>(4.7kb upstream of HLA-DQA1) | $3.15 \times 10^{-89}$ | 1.77 | 0.01 | 1.04 | 0 |

<sup>a</sup> $r^2$ with rs1233578, the top HLA variant in schizophrenia, was obtained from the GAIN schizophrenia cohort (mgs2); <sup>b</sup>Effect size estimate is for HLA-DPB1\*05:01; <sup>c</sup>Effect size estimate obtained from Asian sample. n.r., not reported; Disease abbreviations as defined in **Table 1**. Bold font indicates statistically significant association with schizophrenia.

The number of genome-wide significant non-HLA risk loci for each of the 19 immune diseases varied from three (NAR) to 144 (IBD). Several variants were associated with more than one immune disease. In total we identified 581 unique variants (563 non-HLA variants and 18 HLA variants) associated with any immune disease at genome-wide significance. We refer to these variants as **immune risk variants**.

### Identifying pleiotropic variants implicated in both immune disease and schizophrenia

First, we evaluated whether there was any evidence of overall risk allele sharing between each of the 19 immune diseases and schizophrenia using a binomial sign test. To do this, we used previously published findings from a GWAS conducted by the Schizophrenia Working Group of the Psychiatric Genomics Consortium [1, 11]. This GWAS represented a meta-analysis of 52 cohorts, comprising a total of 35,476 cases and 46,839 controls, and the full dataset is referred to here as the **PGC2 study**. Overall, the direction of effect for the sets of non-HLA SNPs associated with each of the 19 immune diseases at genome-wide significance was not shared with schizophrenia more than expected by chance (all binomial sign test p>0.05, **S1 Fig**). Thus, we did not observe evidence of risk allele sharing between any immune disease and schizophrenia when using a stringent genome-wide significance threshold to define immune risk variants. We also evaluated the collective association of 261 LD-independent, non-HLA immune risk variants associated with at least one of the 19 immune-mediated diseases, for which linkage disequilibrium (LD) Score and minor allele frequency (MAF) information were available in the European LD Score database [16]. We found significant deviation from the theoretical null in schizophrenia for immune risk SNPs (λ=1.46). However, when we compared the association of immune risk SNPs to that of similar randomly selected SNP sets (**Supplementary Methods**) we observed no evidence of enrichment (**S2 Fig**, p=0.66), indicating that immune risk SNPs were not associated with schizophrenia more than expected by chance given the polygenic nature of schizophrenia.

Next, we identified potential pleiotropic variants by evaluating the association of individual immune risk variants with schizophrenia. We considered SNPs associated with schizophrenia at p<8.6×10^−5^ (Bonferroni correction for 581 tests, 563 non-HLA and 18 HLA variants) to have pleiotropic effects. Given the size of the schizophrenia GWAS, we had over 80% power to detect pleiotropic SNPs assuming an OR≥1.12 in schizophrenia.

Within the MHC region, we observed four HLA risk alleles associated with both immune disease and schizophrenia, particularly in the class II HLA region (**Table 2, S3 Fig**). These HLA risk alleles were the strongest MHC region associations for AA (HLA-DRB1 #37 Asn), CEL (HLA-DQB1 #74 Ala), PSC (HLA-B*08:01), and SJO (HLA-DQB1*02:01). The presence of HLA-DRB1 #37 Asn conferred a protective association in both AA and schizophrenia, but the remaining HLA variants showed the opposite direction of effect in schizophrenia compared to immune disease (**Table 2**, **S3 Fig**). Notably, none of these four HLA variants were significantly associated with schizophrenia in previous conditional analyses [9, 11], suggesting that their association with schizophrenia may be driven by LD with other causal variants in the region rather than true pleiotropy. Thus, we did not focus additional analyses on these variants.

Outside of the MHC region, five immune risk variants showed potential pleiotropic effects, with the risk allele for immune disease also conferring risk for schizophrenia. These variants have been previously implicated in CRO (rs6738825, rs13126505, rs1734907 [34, 35]), MS (rs7132277 [36]), and CEL (rs296547 [37]). To evaluate the pleiotropic potential of these non-HLA variants, we used conditional and joint analysis (COJO) [38] to perform association analyses in the PGC2 schizophrenia GWAS conditioning on each of the five immune risk variants (**S4 Fig**). In the setting of true pleiotropy, no significant associations should remain after conditioning on the immune risk variants (statistically, all p>8.6×10^−5^). Consistent with pleiotropy, we observed no remaining associations with schizophrenia after conditioning on rs296547 (top SNP after conditioning: rs111530734, p=1.19×10^−3^), rs1734907 (top SNP after conditioning: rs11768688, p=9.79×10^−4^), and rs13126505 (top SNP after conditioning: rs112786981, p=4.58×10^−4^). Significant associations with schizophrenia remained after conditioning on rs6738825 (top SNP after conditioning: rs111744017, p=8.03×10^−6^) and rs7132277 (top SNP after conditioning: rs74240770, p=1.37×10^−8^), suggesting there were independent causal variants driving the associations in these regions for schizophrenia and immune disorders.

In order to prioritize genes underlying the identified pleiotropic SNPs (rs296547, rs1734907, rs13126505), we performed an integrative analysis of GWAS summary statistics with methylation quantitative trait loci (mQTL) and expression quantitative trait loci (eQTL) studies using SMR and HEIDI [39, 40] (**Materials and Methods**). Notably, rs296547 was not genotyped in the eQTL dataset, and we used rs404339 as a proxy SNP (r^2^=0.85 in 1000 Genomes Phase 3 CEU Population [41]) in SMR analyses of gene expression analyses for rs296547. We observed that rs1734907 was an mQTL (β = 0.47, P = 2.13×10^−26^) and eQTL (β = −0.24, P = 3.54×10^−10^) for *EPHB4* in peripheral blood (**S2 Table, Fig 1**). Furthermore, we observed consistent pleiotropic associations for rs1734907 with schizophrenia and *EPHB4* DNAm (β_SMR_ = −0.14, P_SMR_ = 3.58×10^−5^, P_HEIDI_ = 0.12), schizophrenia and *EPHB4* expression (β_SMR_ = −0.28, P_SMR_ = 2.63×10^−4^, P_HEIDI_ = 0.17), and *EPHB4* DNAm and *EPHB4* expression (β_SMR_ = 1.98, P_SMR_ = 6.56×10^−8^, P_HEIDI_ = 0.011). Thus, there was consistent association across molecular phenotypes and schizophrenia at the *EPHB4* locus, suggesting this gene may be driving the association of rs1734907 in schizophrenia (**Fig 1**). Notably,*TRIP6* is also a candidate functional gene underlying the association of rs1734907 with schizophrenia. We observed pleiotropic association for rs1734907 with schizophrenia and *TRIP6* DNAm with inconsistent direction of effect (β_SMR_ = 0.15, P_SMR_ = 5.00×10^−5^, P_HEIDI_ = 0.17 for probe cg18683606; β_SMR_ = - 0.12, P_SMR_ = 2.32×10^−5^, P_HEIDI_ = 0.18 for probe cg27396824), a trend for association with schizophrenia and *TRIP6* expression (β_SMR_ = −0.33, P_SMR_ = 6.38×10^−4^, P_HEIDI_ = 0.14), but no significant association with *TRIP6* DNAm and *TRIP6* expression. The other pleiotropic SNPs (rs296547, rs13126505) did not demonstrate consistent localization to a particular gene across traits and molecular phenotypes (**Table 3, S2 Table**). We observed that rs296547 was an mQTL for *C1orf106* (β = −1.04, P < 10^−30^), and found pleiotropic associations for rs296547 with schizophrenia and *C1orf106* DNAm but no other phenotypes (**Table 3, S2 Table**). Similarly, we observed that rs13126505 was an mQTL (β = 0.49, P = 4.03×10^−16^) and eQTL (β = −0.27, P = 3.54×10^−10^) for S*LC39A8*, and found pleiotropic associations for rs13126505 with schizophrenia and *SLC39A8* DNAm along with schizophrenia and *SLC39A8* expression, but not *SLC39A8* DNAm and expression (**Table 3, S2 Table**).

**Table 3.** Immune disease risk SNPs showing pleiotropic effect in schizophrenia

| SNP (chr:bp) | Immune Disease | Risk Allele/<br>Non-Risk Allele | Immune OR (95% CI); p<sup>a</sup> | Schizophrenia OR (95% CI); p | Nearby Genes | eQTL<sup>b</sup> | mQTL<sup>c</sup> | Genomic associations co-localizing to this gene<sup>d</sup> |
| --- | --- | --- | --- | --- | --- | --- | --- | --- |
| rs296547<sup>e</sup><br>(chr1:200892137) | CEL [37] | G/A | 1.12<br>(1.09-1.16);<br>4.11x10<sup>-9</sup> | 1.04<br>(1.02-1.07);<br>6.17x10<sup>-5</sup> | *CAMSAP2*<br>*C1orf106*<br>*KIF21B*<br>*CACNA1S*<br>*ASCL5* | n.s. | *C1orf106*,<br>decreased<br>methylation | SCZ-mQTL |
| rs13126505<br>(chr4:102865304) | CRO<sup>f</sup> [35] | A/G | 1.17<br>(1.10-1.25);<br>2.33x10<sup>-10</sup> | 1.14<br>(1.10-1.19);<br>1.19x10<sup>-8</sup> | *BANK1*<br>*SLC39A8*<br>*NFKB1* | *SLC39A8*,<br>decreased<br>expression | *SLC39A8*,<br>increased<br>methylation | SCZ-eQTL,<br>SCZ-mQTL |
| rs1734907<br>(chr7:100315517) | CRO<sup>f</sup> [35] | A/G | 1.16<br>(1.11-1.21);<br>1.67x10<sup>-13</sup> | 1.07<br>(1.04-1.10);<br>7.55x10<sup>-6</sup> | *TFR2*<br>*ACTL6B*<br>*GNB2*<br>*GIGYF1*<br>*POP7*<br>*EPO*<br>*ZAN*<br>*EPHB4*<br>*SLC12A9* | *EPHB4*,<br>decreased<br>expression<br>*TRIP6*,<br>decreased<br>expression | *EPHB4*,<br>increased<br>methylation<br>*TRIP6*,<br>inconsistent<br>effect<br>across<br>probes | SCZ-eQTL,<br>SCZ-mQTL,<br>eQTL-mQTL<br>SCZ-eQTL,<br>eQTL-mQTL |

<sup>a</sup>Effect sizes and p-values reported based on Immunobase curation, which reports statistics from meta-analysis of discovery and replication datasets where available; <sup>b</sup>eQTL data was obtained from the CAGE study [82] which measured gene expression in peripheral blood. Effect on expression (increased/decreased) corresponds to the risk allele; <sup>c</sup>mQTL data was obtained from a meta-analysis of the Brisbane Systems Genetics Study [80] and Lothian Birth Cohorts of 1921 and 1936 [81], which measured DNA methylation in peripheral blood. Effect on expression (increased/decreased) corresponds to the risk allele; <sup>d</sup>Significant SMR and HEIDI [39, 40] results indicating co-localization of genomic associations with the gene of interest in schizophrenia-eQTL (SCZ-eQTL), schizophrenia-mQTL (SCZ-mQTL), and eQTL-mQTL (eQTL-mQTL) datasets; <sup>e</sup>eQTL data were unavailable for rs296547, and rs404339 was used as a proxy SNP ( $r^2=0.85$ in 1000 Genomes Phase 3 CEU Population [41]); <sup>f</sup>Also associated with inflammatory bowel disease; n.s., no statistically significant findings; Disease abbreviations as defined in **Table 1**.

**Fig 1.**
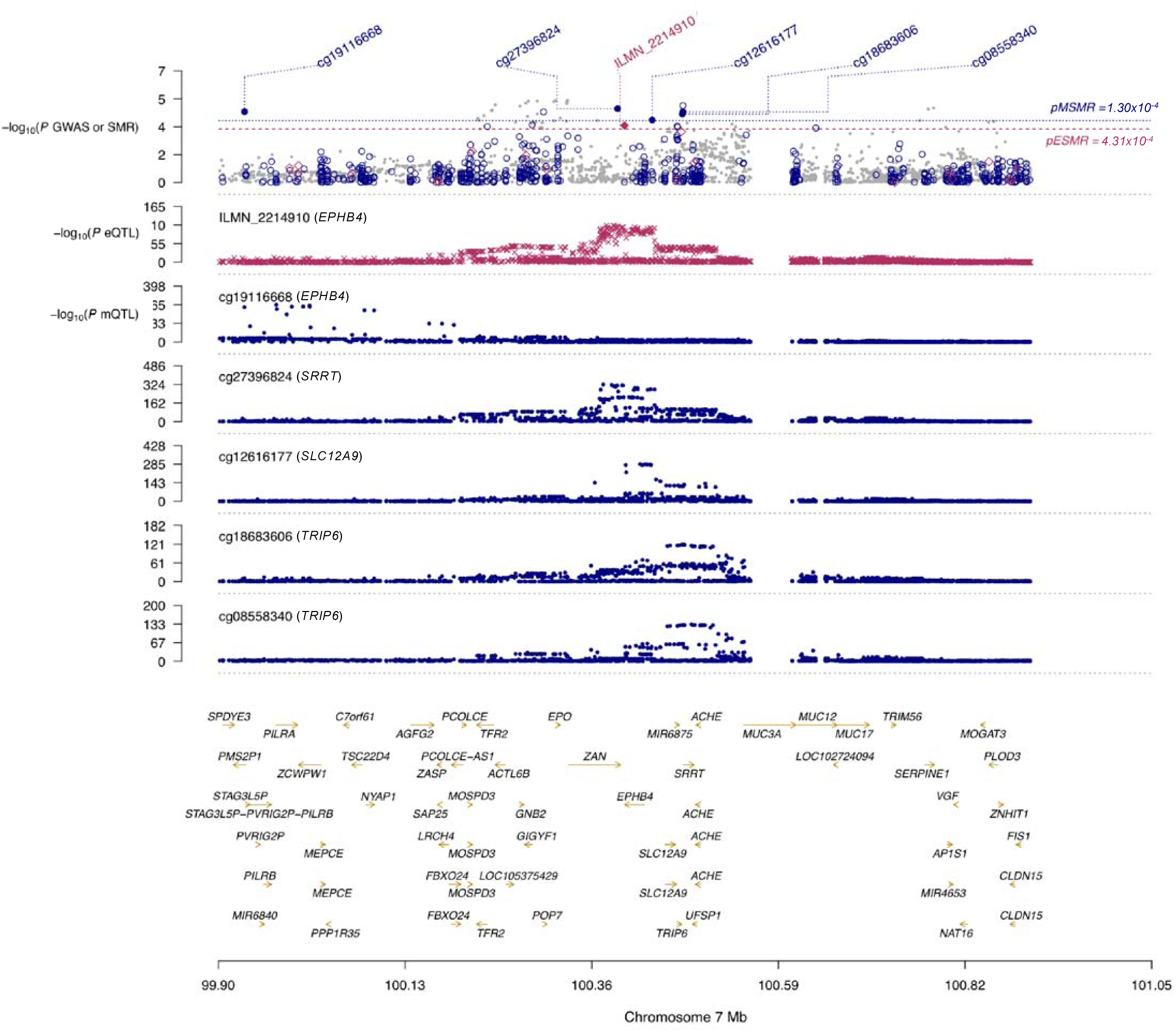
Prioritizing genes driving the pleiotropic association of rs1734907 in Crohn’s disease and schizophrenia. Associations for SNP and SMR analyses across GWAS, eQTL, and mQTL datasets. Top plot gray circles illustrate SNP association (-log_10_ p-value) with schizophrenia in the PGC-2 GWAS, while pink diamonds and blue circles indicate results of SMR tests (-log_10_ p-value) for association of gene expression and DNAm with schizophrenia, respectively, with solid shading indicating probes passing the HEIDI test. Middle plot illustrates SNP association (-log_10_ p-value) with gene expression from peripheral blood eQTL dataset. Lower plots illustrate SNP association (-log_10_ p-value) with gene methylation from peripheral blood mQTL dataset.

### Detecting genetic correlations between immune disease and schizophrenia

Our immune risk variant set captured only those variants associated with immune diseases at genome-wide significance in current GWASs. Given the polygenicity of immune-related diseases, there are 100s to 1,000s of additional variants associated with each disease which have not yet been identified [42]. To evaluate sharing of risk alleles between immune diseases and schizophrenia using a broader set of variants, we used PRS [30, 32] and LDSC [16].

For each of the 14 immune diseases with available genome-wide summary statistics, we constructed genetic risk scores (GRSs) at a range of p-value thresholds (p_T_) as in previous studies [12], and tested for the association of these GRSs with schizophrenia in a refined subset of the PGC2 study (17,000 cases and 20,655 controls) which excluded samples shared with the immune disease GWASs. To benchmark our findings in immune diseases, we also analyzed human height [43] and included previously published PRS results for bipolar disorder [12]. We considered immune diseases with PRS p<0.002 at any p_T_ to show significant genetic overlap with schizophrenia (Bonferroni correction for 14 immune diseases tested in both sexes, 0.05/(14*2) ≈ 0.002). Commonly used goodness-of-fit estimates obtained from PRS (such as *β*_GRS_ and Nagelkerke’s pseudo-R^2^) lack meaningful interpretation, which makes it difficult to compare these estimates across studies [44]. For these reasons we chose to interpret the direction of effect (i.e. positive or negative correlation) obtained from *β*_GRS_, but not to interpret or compare the degree of genetic sharing between immune diseases and schizophrenia. For further details of our PRS approach, see **Materials and Methods**. Using PRS, we had over 80% power to detect genetic covariance with schizophrenia ranging from 0.02 to 0.03 for most of the immune diseases, although some showed less than 80% power in this range (PSO, SLE, VIT; **S5 Fig**).

As previously described, bipolar disorder PRSs were significantly associated with schizophrenia (p<1×10^−50^ at p_T_<1) [12]. Surprisingly, human height PRSs were also significantly associated with schizophrenia (p=1×10^−11^ at p_T_<1, **S3 Table**). Height was analyzed as a negative control based on its previously reported lack of genetic correlation with schizophrenia using LDSC [16]. Using PRS, we observed that genetic liability for increased height protected against schizophrenia (β_GRS_=-0.11 at p_T_<1). The significant inverse association of height PRSs with schizophrenia case-status we observed may reflect the greater sensitivity of this approach to subtle population stratification, sample sharing, and/or true genetic overlap.

Genetic scores including the HLA region were significant for CEL, NAR, PBC, PSO, RA, SLE, SSC, T1D, and UC (p<0.002 at multiple p_T_, **S4 Table**). Height was not included in these analyses, given that HLA variants have not been associated with height in previous GWAS [43]. With the exception of CEL (β_GRS_ ≈ 0.04 at p_T_<5×10^−8^, 1×10^−4^, and 1×10^−3^), all immune diseases exhibited a positively associated PRS with schizophrenia case-status (all β_GRS_>0, **S4 Table**). For CEL, RA, SLE, and SSC only those PRSs constructed using the most stringent p-value cutoffs (5×10^−8^, 1×10^−4^, 1×10^−3^) were significantly associated with schizophrenia. To evaluate whether the HLA region alone was driving the observed genetic sharing, we constructed PRSs excluding this region. After excluding HLA variants, genetic scores for NAR, PBC, PSO, SLE, T1D, and UC remained significantly associated with schizophrenia (**Table 4, S6 Fig**). Because the genetic overlap between these six immune diseases and schizophrenia was not driven by a single HLA variant of large effect, we focused on these findings for the remainder of our analyses.

**Table 4.** Estimated phenotypic and genome-wide genetic correlations between schizophrenia and other traits

| Trait | $h^2 \pm SE^a$ | $r_p$ | PRS | | | | LDSC | |
| --- | --- | --- | --- | --- | --- | --- | --- | --- |
| | | | best $p_T$ | $\beta_{GRS} \pm SE$ | $R^2(\%)$ | p | $r_g \pm SE$ | p |
| BPD (+)<sup>b</sup> | 0.26 ± 0.01 |  | 1 | n.a. | 2.1 | <10<sup>-50</sup> | 0.75 ± 0.05 | 4.02x10<sup>-57</sup> |
| HGT (-) | 0.34 ± 0.19 |  | 1 | -0.11 ± 0.02 | 0.064 | 1.22x10<sup>-11</sup> | 7.47x10<sup>-5</sup> ± 0.02 | 0.99 |
| NAR | 0.31 ± 0.09 | n.a. | 1 | 0.04 ± 0.01 | 0.017 | 4.07x10<sup>-4</sup> | 0.213 ± 0.10 | 0.03 |
| PBC | 0.46 ± 0.08 | 0.11 | 0.3 | 0.07 ± 0.01 | 0.053 | 8.05x10<sup>-10</sup> | 0.131 ± 0.05 | 4.00x10<sup>-3</sup> |
| PSO | 0.27 ± 0.09 | 0.13 | 0.3 | 0.04 ± 0.01 | 0.025 | 2.26x10<sup>-5</sup> | 0.182 ± 0.07 | 7.80x10<sup>-3</sup> |
| SLE | 0.15 ± 0.02 | 0.05 | 0.5 | 0.07 ± 0.01 | 0.047 | 1.50x10<sup>-8</sup> | 0.127 ± 0.045 | 4.60x10<sup>-3</sup> |
| UC | 0.23 ± 0.03 | -0.001 | 0.4 | 0.04 ± 0.01 | 0.018 | 3.74x10<sup>-4</sup> | 0.106 ± 0.04 | 4.00x10<sup>-3</sup> |

$R^2$ and $h^2$ are reported on the liability scale for all diseases; <sup>a</sup> $h^2$ was estimated using LDSC; <sup>b</sup>results reported are from previously published analyses by the Cross-Disorder Working Group of the Psychiatric Genomics Consortium [12]; (+), positive control; (-), negative control; n.a., not available; SE, standard error; $r_g$ , genetic correlation; $r_p$ , expected phenotypic correlation based on epidemiological data (see **Materials and** **Methods** for details of $r_p$ estimation).

Given the potential sensitivity of PRS to artificial genetic overlap highlighted in our analysis of height, we wanted to assess whether cryptic sample sharing between the immune and schizophrenia GWASs could be driving the shared genetic liability that we observed. To do this, we conducted leave-half-out analyses. If the observed genetic overlap was driven by samples shared between certain schizophrenia cohorts and the immune disease GWASs, the GRS association should not be consistently observed across subsamples leaving out half of the schizophrenia cohorts. Across 1,000 subsamples (N_cases_ ranging from 3,985-13,074) leaving out a randomly selected 14 cohorts, we observed a high proportion of subsamples with GRSs significantly associated with schizophrenia (p<0.05 at p_T_<1) for height (0.99), NAR (0.72), PBC (0.95), PSO (0.84), SLE (0.97), T1D (0.95), and UC (0.70) suggesting our findings were not driven by sample sharing.

To further validate our finding of genetic overlap between schizophrenia and these six immune-mediated diseases using PRS, we applied an independent method (LDSC) for estimating genome-wide genetic correlation between traits that is robust to sample sharing [16]. For LDSC analyses, we used summary statistics from the 49 European-ancestry cohorts in the PGC2 study (31,335 cases and 38,765 controls) [1]. Unlike PRS, LDSC provides an interpretable and comparable estimation of genetic sharing between two traits in the form of genetic correlation (r_g_) values. Notably, LDSC is less sensitive than PRS and is not robust when applied to genetic data obtained from specialty chips (e.g. Immunochip) [16]. We did not carry T1D forward for LDSC analysis, due to failure of this dataset on quality control measures (liability scale h^2^ >1, likely secondary to inflated effective sample size due to genotyping on Immunochip). Given that this was a secondary analysis, we considered immune diseases with r_g_ p<0.05 to show significant genetic overlap with schizophrenia.

As previously reported [16], our positive control (bipolar disorder) showed significant genetic overlap with schizophrenia (r_g_=0.75±0.05, p=8.5×10^−60^; **Fig 2, Table 4**). In contrast to our PRS results, but in agreement with previous findings [16], our negative control (height) showed no such overlap using LDSC (r_g_=-0.004±0.02, p=0.84; **Fig 2**, **Table 4**). With respect to immune diseases, LDSC confirmed significant genetic overlap with schizophrenia for PBC, PSO, SLE, and UC (r_g_=0.10-0.18, **Fig 2**, **Table 4**) indicating the association of GRSs for these diseases was not driven by shared samples. Notably, genetic correlations for PSO and SLE did not survive correction for the 14 tests performed (**Table 4**). We also observed significant genetic overlap with schizophrenia for NAR using LDSC, with the caveat that this dataset was genotyped using Immunochip and did not survive multiple testing correction (**Fig 2**, **Table 4**). Overall, LDSC provided consistent results for the immune diseases showing significant genetic sharing with schizophrenia by PRS.

**Fig 2.**
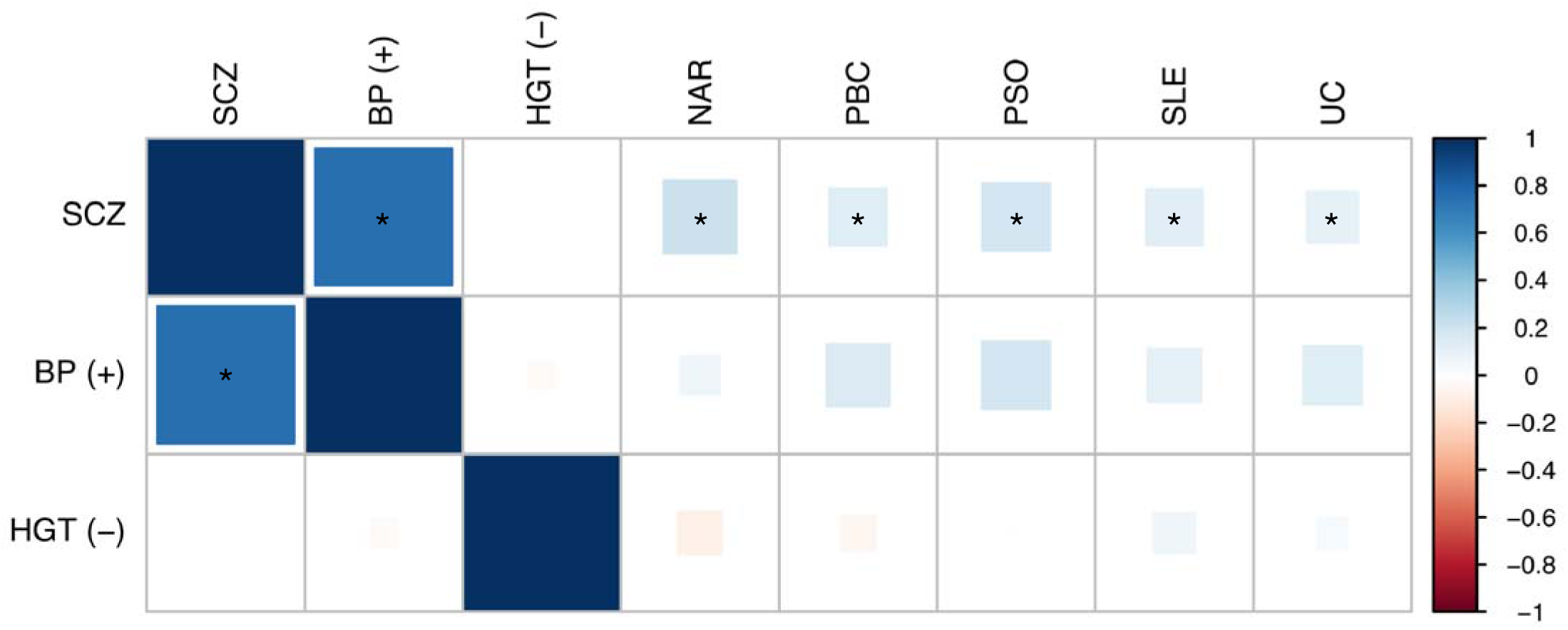
Genetic correlation between schizophrenia and other traits. Genetic correlation between schizophrenia, bipolar disorder, height, and 14 immune diseases was estimated using cross-trait LDSC [16]. Colour of square indicates strength of genetic correlation (red, negative correlation; blue, positive correlation). Size of square indicates statistical significance (larger, more significant *p*-value). Asterisks indicate genetic correlations that are statistically significant at *p* < 0.05 threshold. BP, bipolar disorder; CEL, celiac disease; CRO, Crohn’s disease; HGT, height; IBD, inflammatory bowel disease; JIA, juvenile idiopathic arthritis; MS, multiple sclerosis; NAR, narcolepsy; PBC, primary biliary cirrhosis; PSO, psoriasis, RA, rheumatoid arthritis; SLE, systemic lupus erythematosus; SSC, systemic sclerosis; T1D, type 1 diabetes; UC, ulcerative colitis; VIT, vitiligo.

### Benchmarking genetic correlations between immune disease and schizophrenia with epidemiological data

To determine how much of the phenotypic correlation between schizophrenia and immune-mediated diseases was explained by the genetic correlations we observed, we benchmarked significant genetic correlations between schizophrenia and immune-mediated disorders relative to the expected phenotypic correlations from epidemiological data (**Materials and Methods**). Using incidence of immune diseases in schizophrenia reported in a large population-based study [3], we estimated phenotypic correlations between schizophrenia and PBC, PSO, SLE, and UC. We were unable to estimate phenotypic correlation for NAR and schizophrenia, given that there were no estimates in the literature of the incidence of NAR in schizophrenia. For PBC, PSO, and SLE we observed small positive genetic correlations with schizophrenia that were consistent with the epidemiological data (PBC: r_g_ = 0.131 ± 0.05, r_p_ = 0.112; PSO: r_g_ = 0.182 ± 0.07, r_p_ = 0.130; SLE: r_g_ = 0.130 ± 0.05, r_p_ = 0.048). For UC we observed a small positive estimate of genetic correlation (r_g_ = 0.106 ± 0.04) while there was no strong evidence for any correlation between UC and schizophrenia in the epidemiological data (r_p_ = −0.001). Importantly, while the MHC region contains risk factors for both schizophrenia and immune-mediated diseases, our genetic correlation estimates were obtained considering only SNPs outside of the MHC region due to unusual LD in this region [45].

### Exploring sex-dependent genetic correlations between immune disease and schizophrenia

Given the significant sex bias of autoimmune diseases, with women at greater risk overall [46], we hypothesized that there may be sex-dependent genetic overlap between schizophrenia and some immune-mediated diseases. We therefore performed sex-stratified PRS, testing the association of height and immune disease GRSs with schizophrenia separately in males and females of the PGC2 study. Genetic scores for height showed significant association with schizophrenia in both males and females. Three of the immune diseases (PBC, PSO, T1D) with significant main effects showed sex-dependent effects, with greater signal among males (**S5 Table**). Additionally, although genetic scores for MS were not significantly associated with schizophrenia in the total sample there was significant association among males (R^2^=0.03, p=1.26×10^−3^ at p_T_<1; **S5 Table**).

Given the greater statistical power for the male subset of the schizophrenia GWAS, we performed simulations by selecting random subsamples of male cases and controls equal in size to the female sample (5,321 cases and 9,094 controls). If the stronger genetic overlap between schizophrenia and MS, PBC, PSO, and T1D among males was driven by the larger sample size rather than a true sex-dependent effect, there should be no consistent association of GRSs with schizophrenia in these subsamples. Across 1,000 subsamples, the proportion with significant GRSs (p<0.002 at p_T_<1) was high for PBC (0.94) and T1D (0.87), suggesting our finding of a greater pleiotropic effect among males for these diseases was not driven solely by lower statistical power among females; this was not the case for PSO (0.59) or MS (0.21).

Next, we performed formal statistical tests for an interaction between sex and genetic scores for these four immune diseases. We observed a nominally significant interaction for MS (p<0.05 at several p_T_; **S5 Table**), noting that this finding did not survive correction for multiple testing. The remaining immune diseases did not show significant sex interactions, although the direction of effect was consistent with a greater pleiotropic effect in males (**S5 Table**).

## Discussion

Using a variety of statistical approaches, we provide evidence of shared genetic risk for schizophrenia and several immune diseases. Within the MHC region, we identified four HLA variants showing statistically significant association with schizophrenia. An important caveat is that these four variants were not the top variants in their respective regions of association with schizophrenia, and were not primary drivers of the MHC association in schizophrenia in stepwise conditional analyses [9]. Therefore, the biological significance of these particular HLA variants in schizophrenia is likely limited.

Outside of the MHC region, we identified three SNPs with pleiotropic effects - influencing risk for both celiac disease (CEL) (rs296547) or Crohn’s disease (CRO) (rs1734907, rs13126505) and schizophrenia. Integration of GWAS, mQTL, and eQTL data implicated *C1orf106, SLC39A8*, and *EPHB4* or *TRIP6* as functional candidates driving the pleiotropic association of rs296547, rs13126505, and rs1734907, respectively. Overall, our findings provide the strongest evidence for a model in which genetic variation at rs1734907 (∼85kb upstream of *EPHB4*) increases DNA methylation, upregulates *EPHB4* expression, and decreases the risk of schizophrenia. While DNA methylation is classically associated with gene silencing, the effect of methylation on transcription depends on the genomic context [47]; for instance, methylation of silencers or insulators eliminates transcription-blocking activity thereby promoting gene expression [48, 49]. *EPHB4* is a transmembrane tyrosine kinase receptor that coordinates cell movement via bidirectional intercellular signaling at sites of direct cell-to-cell contact [50]. In the brain, ephrin signaling mediates synaptic plasticity by initiating and stabilizing neuronal synapse formation (reviewed by [51]). An analogous role has not yet been discovered in the immune system, possibly due to the much shorter lifespan of immunological synapses between lymphocytes and antigen presenting cells (minutes) as compared to neuronal synapses (years) [52, 53]. Interestingly, ephrin signaling attenuates the migration responses of both neurons and immune cells toward chemoattractants *in vitro* [54, 55]. Thus, disrupted pathfinding may be a shared risk mechanism by which *EPHB4* contributes to immune disease and schizophrenia. The hypotheses raised by our findings require further validation. If the association of rs1734907 with CRO and schizophrenia is robustly replicated in future GWASs, functional studies will be needed to investigate both the genetic mechanism by which rs1734907 (or a causal variant in LD with this SNP) influences *EPHB4* transcription, and the biological mechanism by which increased *EPHB4* expression influences susceptibility to CRO and schizophrenia. With the multi-kinase inhibitor dasatinib already on the market for treatment of chronic myeloid leukemia [56] and other EphB4 inhibitors currently in Phase II trials [57–60], the potential for future drug repurposing makes *EPHB4* an attractive candidate for further investigation.

We observed genome-wide sharing of risk variants for schizophrenia and six immune diseases (narcolepsy (NAR), primary biliary cirrhosis (PBC), psoriasis (PSO), systemic lupus erythematosus (SLE), type 1 diabetes (T1D), and ulcerative colitis (UC)) using PRS, all of which have been previously reported to co-occur with schizophrenia in epidemiological studies [3, 5, 61]. The strongest evidence of shared genetic risk emerged for PBC, PSO, SLE, and UC, which also showed robust genetic correlation with schizophrenia using LDSC. With the exception of UC, the small positive genetic correlations observed between these immune diseases and schizophrenia (r_g_ ∼ 0.1) were consistent with phenotypic correlations observed in epidemiological data. Thus, currently available genetic data suggest that shared genetic risk contributes to the co-occurrence of PBC, PSO, and SLE in schizophrenia. Possible explanations for this sharing of genetic risk include the presence of a subgroup of “autoimmune-like” schizophrenia cases and/or sharing of specific biological pathways between schizophrenia and these particular immune diseases.

To our knowledge, this is the first time that sex-dependent genetic correlation with immune diseases has been investigated in schizophrenia. We found nominal evidence of male-specific genetic correlation for multiple sclerosis (MS), and a stronger pleiotropic effect among males for PBC, PSO, and T1D although the latter were not statistically significant. Interestingly, animal studies indicate that sex hormones have opposing effects on predisposition to schizophrenia and autoimmunity; estrogen has been reported to protect against the development of schizophrenia [62], while androgens appear to protect against the development autoimmune diseases [63, 64]. We emphasize that our sex-dependent findings require validation in independent samples. If replicated, one possibility is that sex hormones modulate pathogenesis among genetically vulnerable individuals, making males more likely to develop schizophrenia and females more likely to develop autoimmune diseases.

Our work was subject to several important limitations. Firstly, genome-wide summary statistics were not available for all of the immune diseases, resulting in a more limited analysis of 14 diseases. For five of these diseases (CEL, juvenile idiopathic arthritis (JIA), MS, NAR, T1D) summary statistics were obtained from Immunochip rather than GWAS, providing incomplete coverage of the genome for comparison with schizophrenia and biasing the genetic correlation estimates obtained by LDSC. Secondly, GRSs for human height – analyzed as a negative control – showed stronger association with schizophrenia than any of the immune diseases. An inverse epidemiological relationship between height and schizophrenia has been reported [65, 66], consistent with our PRS findings. The reasons for the discrepancy between PRS and LDSC, which showed no genetic correlation between height and schizophrenia (as previously reported [16]) are unclear. One explanation is that PRS, which uses individual-level genotype data as opposed to summary statistics, is a more sensitive method to detect true genome-wide sharing of risk alleles. If this is the case, it raises a broader question regarding how much genetic overlap is expected across complex traits in general using the PRS approach. Recent work suggests that pleiotropy is pervasive across human diseases, and that this phenomenon is driven at least in part by the polygenic nature of complex traits [21]. If this is the case, the extreme polygenicity of human height (more than 100,000 common variants estimated to exert independent causal effects [67]) may be driving the pleiotropy we observed between height and schizophrenia using PRS. An alternative explanation that must be considered is that PRS may be more vulnerable to confounding by cryptic population stratification, LD, or sample sharing.

Despite these limitations, our work adds to a growing body of evidence suggesting that schizophrenia and immune diseases share genetic risk factors. There are conflicting reports in the literature with respect to the specific immune diseases demonstrating genetic overlap with schizophrenia, and the direction of effect (positive or negative genetic correlation). Genetic overlap with schizophrenia has been previously investigated for nine of the 19 immune diseases studied here. Genome-wide genetic correlation with schizophrenia has been previously reported for CRO [23–25, 27], MS [28], PBC [25], PSO [25, 29], rheumatoid arthritis (RA, both positive [23, 24] and negative [31] genetic correlations), SLE [24, 25], T1D [23], and UC [24–27] (see **S1 Table** for a summary of previous studies). Our results are consistent with previously reported genetic overlap between schizophrenia and PBC [25], PSO [25], SLE [24, 25], T1D [23], and UC [24, 25]. While we did not observe genetic correlation between schizophrenia and MS in the total sample, there was a significant sex-dependent effect with genetic correlation observed among males. We provide new evidence of genetic correlation with NAR (not previously investigated). Notably, we did not find any significant genetic correlation between schizophrenia and RA. Despite the robust inverse epidemiological association between schizophrenia and RA [8], the genetic association is less consistent. Using methods based on summary statistics (including PRS and LDSC), four previous studies reported no evidence of pleiotropy between schizophrenia and RA [8, 16, 25, 30], while two studies reported positive genetic correlation [23, 24]. Notably, Lee *et al.* reported an inverse genetic correlation – in keeping with the observed epidemiological effect – using restricted maximum likelihood (GREML), a method utilizing full genotype data which has greater statistical power to detect small pleiotropic effects than PRS or LDSC [31]. Given the modest and potentially sex-dependent genetic correlations observed in the present study, subtle differences in statistical power across studies using different statistical methods and GWAS datasets may explain these discrepant findings. As genetic samples continue to grow, and our understanding of the degree of genetic overlap expected among complex traits evolves, it will be worthwhile to revisit these analyses.

Overall, our analyses provide statistical evidence supporting extensive pleiotropy between immune diseases and schizophrenia. Our results highlight *EPHB4*, a transmembrane receptor that coordinates cell migration and has dual roles in immune cell and neuronal pathfinding, as a promising candidate for future functional studies. More broadly, our findings indicate that common genetic variants influencing the risk of immune diseases – in particular NAR, PBC, PSO, SLE, and UC – are also involved in schizophrenia. Studies identifying the cell types and biological pathways that may be driving this genetic overlap are needed, and will hopefully provide further insights into the pathophysiology of schizophrenia. In the meantime, our work supports the emerging hypothesis that pathogenic mechanisms are shared across immune and central nervous system disorders.

## Materials and Methods

### Samples and quality control

We used either imputed genotype data or summary statistics generated as described in the original GWASs. For sample details, see **Table 1.**

### Schizophrenia dataset

We used data from the PGC2 study [1]. For analyses of non-HLA genome-wide significant risk variants for immune diseases we used publicly available summary statistics from the total dataset (52 cohorts; 35,476 cases and 46,839 controls) [1]. For PRS analyses we used all 36 European ancestry case-control cohorts with available individual-level genotype data (25,629 cases and 30,976 controls). For analyses including HLA variants we used a further refined 31 European ancestry case-control cohorts (20,253 cases and 25,011 controls) with high-quality coverage of the MHC region, as previously described [11].

### Immune disease datasets

To estimate the extent of genetic overlap between schizophrenia and immune diseases, we obtained full GWAS or Immunochip summary statistics for 14 of the 19 immune diseases (five immune diseases were not included in PRS analyses due to lack of available summary statistics). We obtained publicly available summary statistics for ten immune diseases (see **URLs**): CEL [68], CRO [69], IBD [69], JIA [70], MS [36], NAR [71], RA [72], SLE [73], T1D [74], and UC [69]. For the following four immune diseases, we obtained summary statistics with permission from the authors: PBC [75], PSO [76], SSC [77], and VIT [78].

### Testing the association of genome-wide significant risk alleles for 19 immune diseases in schizophrenia

For each of the 19 immune diseases, we defined risk loci outside of the MHC region (chromosome 6: 25-34 Mb) using curated GWAS results from ImmunoBase (http://www.immunobase.org; accessed 7 June 2015. For details, see **Supplementary Methods**). Notably, the majority of IBD risk variants were also risk variants for CRO and/or UC. Within the MHC region we considered only the most strongly associated HLA variant (including SNPs, imputed HLA amino acid sites, and classical alleles) for each disease based on univariate analysis in previously published studies (see **Table 2**), because multivariate conditional analyses reporting adjusted effect sizes of independent HLA variants were not available for all immune diseases. In total there were 581 unique variants (563 non-HLA variants and 18 HLA variants) associated with any immune disease at genome-wide significance.

First, we tested for shared direction of effect with schizophrenia among SNPs associated with each of the 19 immune diseases using the binomial sign test. Because some immune risk SNPs were associated with multiple diseases with inconsistent direction of effect, we could not evaluate shared direction of effect among the collective set of immune risk SNPs in schizophrenia.

Next, we evaluated the collective association of SNPs associated with any immune disease. First we extracted the p-values for a pruned set of 261 LD-independent, non-HLA immune risk SNPs with linkage disequilibrium (LD) Score and minor allele frequency (MAF) information were available in the European LD Score database [16] from the schizophrenia PGC2 GWAS. We then quantified enrichment of these immune risk SNP associations in schizophrenia using the genomic inflation value λ. We obtained an empirical enrichment p-value by comparing this to λ values from 1,000 equal-sized sets of SNPs drawn from the schizophrenia GWAS summary data, and matched to the immune SNP set for MAF and LD score as these parameters are correlated with GWAS test statistics (see **Supplementary Methods** for details).

Finally, we evaluated the association of each of the 581 variants with schizophrenia using previously published association results for non-HLA [1] and HLA variants [11]. We considered SNPs associated with schizophrenia at p<8.6×10^−5^ (Bonferroni correction for 581 tests, 563 non-HLA and 18 HLA variants) to have pleiotropic effects.

To evaluate the pleiotropic potential of immune risk variants significantly associated with schizophrenia, we performed conditional and joint analysis (COJO) using GCTA [79]. Specifically, we used COJO to perform association analyses in the PGC2 schizophrenia GWAS conditioning on the immune risk variants of interest (i.e. SNPs that were significantly associated with both an immune disease and schizophrenia). In the setting of true pleiotropy, no significant associations with schizophrenia should remain after conditioning on these immune risk variants (statistically, all p>8.6×10^−5^). We used the 1000 Genomes Phase 3 European dataset as a reference panel to calculate LD between SNPs.

To prioritize genes and regulatory elements driving the pleiotropic GWAS loci we identified (associated with both immune disease and schizophrenia, see **Table 3**), we followed the analytic approach described by Wu *et al.* [40]. This approach integrates summary statistics from independent -omics methylation quantitative trait loci (mQTL) studies, expression quantitative trait loci (eQTL) studies, and GWAS to identify SNPs associated with gene expression, DNA methylation, and disease through shared genetic effects.

We obtained mQTL and eQTL data used in Wu *et al.* [40] for genetic regions within a 2Mb window of each pleiotropic SNP. These data and the quality control measures applied have been described in detail elsewhere [40]. Briefly, mQTL summary-level SNP data were from a meta-analysis of the Brisbane Systems Genetics Study [80] and Lothian Birth Cohorts of 1921 and 1936 [81], which comprised 1,980 individuals with DNA methylation measured in peripheral blood. eQTL summary-level SNP data were from the Consortium for the Architecture of Gene Expression (CAGE) study [82], which comprised 2,765 individuals with gene expression levels measured in peripheral blood. GWAS summary-level SNP data for schizophrenia was from the PGC2 study [1].

We applied summary data-based Mendelian randomization (SMR) using GCTA [79] to test for shared associations between the pleiotropic SNPs with DNAm probes and gene expression probes, DNAm probes and schizophrenia, and gene expression probes and schizophrenia. We included DNAm and gene expression probes within 2Mb of the pleiotropic SNPs. We considered significant associations as those with P_SMR_ < 1.30×10^−4^ (0.05/385 tagged genes) for mQTLs and P_SMR_ < 4.31×10^−4^ for eQTLs (0.05/116 tagged genes). Next, we applied the heterogeneity in dependent instruments (HEIDI) test [39] using GCTA [79] to evaluate whether significant shared associations between DNAm, gene expression and schizophrenia were driven by linkage (i.e. separate causal variants in LD exerting genetic effects on DNAm, gene expression, and schizophrenia) or a shared pleiotropic causal variant. We considered genetic effects that passed the HEIDI test (P_HEIDI_ > 0.01) to be driven by a single causal variant. We looked for consistent SMR and HEIDI results across GWAS, mQTL, and eQTL studies to prioritize genes for future functional studies.

### Testing the association of polygenic risk scores for 14 immune diseases in schizophrenia

To evaluate whether common variants influencing risk of immune diseases collectively contribute to schizophrenia, we used PRS [30, 32]. To benchmark the amount of genetic overlap between schizophrenia and immune disease, we included previously published results for bipolar disorder as a positive control [12]. We used human height [43] as a negative control because – despite the inverse epidemiological relationship between height and schizophrenia previously reported [65, 66] – a prior study using cross-trait LDSC reported no genetic correlation with schizophrenia [16].

For 14 immune diseases with available genome-wide summary statistics we performed PRS at a range of p-value thresholds (p_T_) as in previous studies [12]: 5×10^−8^, 1×10^−4^, 1×10^−3^, 0.01, 0.05, 0.1, 0.2, 0.3, 0.4, 0.5, and 1.0 (which included all LD-independent SNPs, **Table 1**). Due to extensive LD in the HLA region, we performed analyses both including the top HLA variant and excluding the HLA region. At each p_T_, we constructed GRSs for each individual *i* in the schizophrenia cohort for each immune disease *h* by calculating the sum of risk-allele dosages (*g*) weighted by their effect sizes (*β*) for that immune disease:

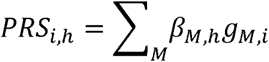

where *M* iterates over all known risk alleles for disease *h, β*_*M,h*_ is the effect size (log odds ratio) of *M* in disease *h*, and *g*_*M,i*_ is the risk-allele dosage of *M* in individual *i*. We then performed logistic regression in R [83] using the stats package [83] to evaluate the association between schizophrenia case-status and GRSs for each immune disease. As in previous studies, statistical significance of the GRSs was estimated based on their logistic regression coefficient [12, 30]. Variance in schizophrenia case-status explained by the GRSs was estimated using the deviation in liability-scale R^2^ between a null model (including 10 ancestry-informative principal components and study site) and the full model (including GRSs in addition to these covariates), calculated as previously described [44] assuming a population prevalence of schizophrenia of 1%. We also estimated Nagelkerke’s pseudo-R^2^ using the fmsb package [84]. We considered immune diseases with GRS p<0.002 at any p_T_ to show significant genetic overlap with schizophrenia (Bonferroni correction for 14 immune diseases tested in both sexes, 0.05/(14*2)=0.002). As in previous studies [12, 30] we did not use Bonferroni correction for the number of p-value thresholds, as these tests are highly correlated.

We excluded eight schizophrenia cohorts using Wellcome Trust Case Control Consortium (WTCCC) controls, due to the use of these samples in the immune disease GWASs. The total schizophrenia sample analyzed by PRS included 37,655 subjects (28 cohorts; 17,000 cases and 20,655 controls). Sex-stratified and formal sex-PRS interaction analyses were performed among the subset of subjects with known sex (9,787 male cases and 9,284 male controls; 5,231 female cases and 9,094 female controls). For details of PRS, see **Supplementary Methods** and **Table 1**.

### Estimating the degree of genetic correlation between schizophrenia and 14 immune diseases

To validate our PRS results and obtain genetic correlation (r_g_) estimates, we performed a secondary analysis using cross-trait LDSC for immune-mediated diseases with significant PRS associations with schizophrenia [16]. Cross-trait LDSC estimates the genetic correlation between two traits using GWAS summary statistics. Similar to the PRS analyses described above, we benchmarked the genetic correlations observed for immune diseases by analyzing bipolar disorder [85] as a positive control and human height [43] as a negative control.

The statistical framework for cross-trait LDSC has been described in detail previously [16]. Briefly, LDSC leverages the relationship between LD and association test statistics to estimate heritability as the slope of the regression of z-scores against LD scores [86]. Cross-trait LDSC is a bivariate extension of this method which estimates genetic covariance as the slope of the regression of the products of z-scores against LD scores using the following equation [16]:

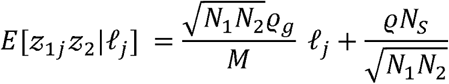

where *z*_*ij*_ denotes the z score for study *i* and SNP*j, 𝓁*_j_ is the LD score [86], *N*_i_ is the sample size for study *i, ϱ*_g_ is the genetic covariance, *M* is the number of SNPs in the reference panel with MAF between 5% and 50%, *N*_S_ is the number of individuals included in both studies, and *ϱ* is the phenotypic correlation among the *N*_S_ overlapping samples. Genetic covariance *ϱ*_g_ is estimated by regressing *z*_1j_ *z*_2_ against 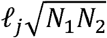, and multiplying the resulting slope by *M*. Statistical significance is assessed using block jackknifing over 200 equally sized blocks of SNPs [16]. Importantly, the MHC region is excluded from LDSC analyses due to its unusual LD structure and genetic architecture [45].

Because LDSC is robust to sample sharing across GWAS [16], we used summary statistics from the 49 European-ancestry cohorts in the PGC2 study (31,335 cases and 38,765 controls) [1]. We used LD Scores from the “eur_w_ld_chr/” files available from https://data.broadinstitute.org/alkesgroup/LDSCORE, computed using 1000 Genomes Project [87] Europeans as a reference panel as previously described [45]. To ensure we were using well-imputed SNPS we filtered all GWAS as previously described [16], including limiting the analysis to HapMap 3 [88] SNPs as implemented in the LDSC script munge_sumstats.py (https://github.com/bulik/ldsc). We estimated liability scale h^2^ for each trait using previously reported prevalence estimates (**S6 Table**), and removed datasets with h^2^>1. Given that this was a secondary analysis, we considered traits with r_g_ p<0.05 to have significant genetic correlation with schizophrenia.

### Benchmarking with epidemiological data

To determine how much of the phenotypic correlation between schizophrenia and immune-mediated diseases was explained by the genetic correlations we observed, we used the approach previously described by Lee *et al.* [31]. Briefly, we benchmarked our significant genetic correlation estimates between schizophrenia and NAR, PBC, PSO, SLE and UC relative to the expected phenotypic correlations from epidemiological data. We obtained estimates of the population risk of schizophrenia (*K*_*SCZ*_), the population risk of each immune disease (*K*_*IMMUNE*_), and the probability of each immune disease among patients with schizophrenia (*K*_*IMMUNE* | *SCZ*_) from the literature as referenced in **S6 Table**. We estimated the phenotypic correlation between schizophrenia and the immune disease of interest (*R*_*SCZ-IMMUNE*_) using the following formula, as derived by Lee *et al.* [31] assuming that the phenotypic liabilities of schizophrenia (*l*_*SCZ*_) and immune disease (*l*_*IMMUNE*_) follow a bivariate normal distribution with mean=0 and standard deviation=1:

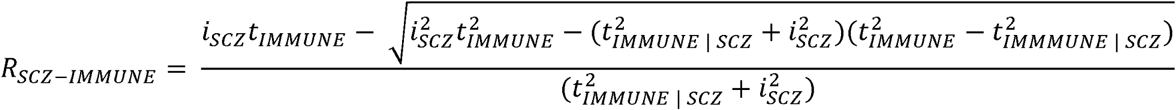

where:

*t*_*SCZ*_ is the liability threshold for schizophrenia:

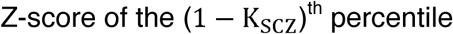

*t*_*IMMUNE*_ is the liability threshold for immune disease:

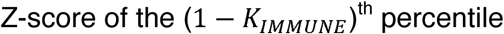

*t*_*IMMUNE* | *SCZ*_ is the liability threshold for immune disease in those with schizophrenia:

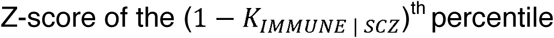

*d*_*SCZ*_ is the “height” of the normal distribution at the schizophrenia liability threshold:

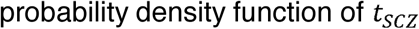

*i*_*SCZ*_ is the mean phenotypic liability of those with schizophrenia:

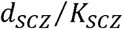

### Statistical power

Power to detect association of individual non-HLA and HLA immune risk variants in schizophrenia was calculated using the Genetic Power Calculator [89] assuming a risk allele frequency (RAF) of 0.05, disease prevalence of 1%, and significance threshold (α) of 8.6×10^−5^. Power for PRS was evaluated using AVENGEME [90, 91], assuming disease and genetic parameters detailed in **S6 Table**.

## URLs

LD Score database: ftp://atguftp.mgh.harvard.edu/brendan/1k_eur_r2_hm3snps_se_weights.RDS

GWAS summary statistics:

- CEL https://www.immunobase.org/downloads/protected_data/iChip_Data/hg19_gwas_ic_cel_tr_ynka_4_19_1.tab.gz
- CRO, IBD, UC ftp://ftp.sanger.ac.uk/pub/consortia/ibdgenetics/iibdgc-trans-ancestry-filtered-summary-stats.tgz
- JIA https://www.immunobase.org/downloads/protected_data/iChip_Data/hg19_gwas_ic_jia_hin_ks_UK_4_19_1.tab.gz
- MS https://www.immunobase.org/downloads/protected_data/GWAS_Data/hg19_gwas_ms_im_sgc_4_19_1.tab.gz
- NAR https://www.immunobase.org/downloads/protected_data/iChip_Data/hg19_gwas_ic_nar_fa_raco_4_19_1.tab.gz
- RA http://www.broadinstitute.org/ftp/pub/rheumatoid_arthritis/Stahl_etal_2010NG/
- SLE https://www.immunobase.org/downloads/protected_data/GWAS_Data/hg19_gwas_sle_be_ntham_4_20_0.tab.gz
- T1D https://www.immunobase.org/downloads/protected_data/iChip_Data/hg19_gwas_ic_t1d_o_nengut_meta_4_19_1.tab.gz

## Supporting information

Pouget2019_SupplementaryInformation

Pouget2019_SupplementaryTables

## Acknowledgements

We thank all study participants, and all research staff at the numerous study sites that made this research possible. Computations were performed on the Genetic Cluster Computer (GCC) hosted by SURFsara (www.surfsara.nl). We thank ImmunoBase for curating genetic risk factors for the immune diseases investigated. We thank the Canadian-US PBC Consortium, the Italian PBC Genetics Study Group and the UK-PBC Consortium for sharing their summary data for the PBC GWAS. We thank the International Inflammatory Bowel Disease Genetics Consortium (IIBDGC) for providing publicly available genome-wide summary statistics for CRO, UC, and IBD. We also thank the International Multiple Sclerosis Genetics Consortium (IMSGC) and the investigators leading the GWASs for CEL, JIA, RA, SLE, and T1D for providing genome-wide summary statistics from their studies on ImmunoBase.

This research was supported in part by a number of funding sources. This research uses resources provided by the Genetic Association Information Network (GAIN), obtained from the database of Genotypes and Phenotypes (dbGaP) found at http://www.ncbi.nlm.nih.gov/gap through dbGaP accession number phs000021.v3.p2; samples and associated phenotype data for this study were provided by the Molecular Genetics of Schizophrenia Collaboration (PI: Pablo V. Gejman, Evanston Northwestern Healthcare (ENH) and Northwestern University, Evanston, IL, USA). JG Pouget is supported by Fulbright Canada, the Weston Foundation, and Brain Canada through the Canada Brain Research Fund - a public-private partnership established by the Government of Canada. B Han is supported by the National Research Foundation of Korea (NRF) (grant 2016R1C1B2013126) and the Bio & Medical Technology Development Program of the NRF (grant 2017M3A9B6061852) funded by the Korean government, Ministry of Science and ICT (MSIT). HM Ollila is supported by the Finnish Cultural Foundation and Academy of Finland (grant 309643). J Martin is supported by grants SAF2015-66761-P from the Spanish Ministry of Economy and Competitiveness and P12-BIO-1395 from from Consejería de Innovación, Ciencia y Tecnología, Junta de Andalucía (Spain). Y Jin, SA Santorico, and R Spritz are supported by the U.S. National Institutes of Health (NIH) grants R01AR045584, R01AR056292, X01HG007484, and P30AR057212 and by institutional research funding IUT20-46 from the Estonian Ministry of Education and Research. MD Mayes is supported by the U.S. NIH grants N01AR02251 and R01AR05528. S Raychaudhuri is supported by the U.S. NIH grants 1R01AR063759, 1R01AR062886, 1UH2AR067677-01, and U19AI111224-01 and Doris Duke Charitable Foundation grant 2013097. J Knight is the Joanne Murphy Professor in Behavioural Science. Funding for the GAIN schizophrenia sample was provided by the U.S. NIH grants R01 MH67257, R01 MH59588, R01 MH59571, R01 MH59565, R01 MH59587, R01 MH60870, R01 MH59566, R01 MH59586, R01 MH61675, R01 MH60879, R01 MH81800, U01 MH46276, U01 MH46289 U01 MH46318, U01 MH79469, and U01 MH79470 and the genotyping of samples was provided through GAIN. The funding sources did not influence the study design, data analysis, or writing of this manuscript.

## References

1. Schizophrenia Working Group of the Psychiatric Genomics Consortium. Biological insights from 108 schizophrenia-associated genetic loci. Nature. 2014;511(7510):421–7.

2. Chan MK, Krebs MO, Cox D, Guest PC, Yolken RH, Rahmoune H, et al. Development of a blood-based molecular biomarker test for identification of schizophrenia before disease onset. Transl Psychiatry. 2015;5:e601.

3. Benros ME, Nielsen PR, Nordentoft M, Eaton WW, Dalton SO, Mortensen PB. Autoimmune diseases and severe infections as risk factors for schizophrenia: a 30-year population-based register study. Am J Psychiatry. 2011;168(12):1303–10.

4. Chen SJ, Chao YL, Chen CY, Chang CM, Wu EC, Wu CS, et al. Prevalence of autoimmune diseases in in-patients with schizophrenia: nationwide population-based study. Br J Psychiatry. 2012;200(5):374–80.

5. Benros ME, Pedersen MG, Rasmussen H, Eaton WW, Nordentoft M, Mortensen PB. A nationwide study on the risk of autoimmune diseases in individuals with a personal or a family history of schizophrenia and related psychosis. Am J Psychiatry. 2014;171(2):218– 26.

6. Eaton WW, Byrne M, Ewald H, Mors O, Chen CY, Agerbo E, et al. Association of schizophrenia and autoimmune diseases: linkage of Danish national registers. Am J Psychiatry. 2006;163(3):521–8.

7. Eaton WW, Pedersen MG, Nielsen PR, Mortensen PB. Autoimmune diseases, bipolar disorder, and non-affective psychosis. Bipolar Disord. 2010;12(6):638–46.

8. Euesden J, Breen G, Farmer A, McGuffin P, Lewis CM. The relationship between schizophrenia and rheumatoid arthritis revisited: genetic and epidemiological analyses. Am J Med Genet B Neuropsychiatr Genet. 2015;168B(2):81–8.

9. Sekar A, Bialas AR, de Rivera H, Davis A, Hammond TR, Kamitaki N, et al. Schizophrenia risk from complex variation of complement component 4. Nature. 2016;530(7589):177–83.

10. Yih Chen J, Ling Wu Y, Yin Mok M, Jan Wu YJ, Lintner KE, Wang CM, et al. Effects of complement C4 gene copy number variations, size dichotomy, and C4A deficiency on genetic risk and clinical presentation of systemic lupus erythematosus in East Asian populations. Arthritis Rheumatol. 2016;68(6):1442–53.

11. Pouget JG, Goncalves VF, Spain SL, Finucane HK, Raychaudhuri S, Kennedy JL, et al. Genome-wide association studies suggest limited immune gene enrichment in schizophrenia compared to 5 autoimmune diseases. Schizophr Bull. 2016;42(5):1176–84.

12. Cross-Disorder Group of the Psychiatric Genomics Consortium. Identification of risk loci with shared effects on five major psychiatric disorders: a genome-wide analysis. Lancet. 2013;381:1371–9.

13. Cross-Disorder Group of the Psychiatric Genomics Consortium. Genetic relationship between five psychiatric disorders estimated from genome-wide SNPs. Nat Genet. 2013;45(9):984–94.

14. Voight BF, Peloso GM, Orho-Melander M, Frikke-Schmidt R, Barbalic M, Jensen MK, et al. Plasma HDL cholesterol and risk of myocardial infarction: a mendelian randomisation study. Lancet. 2012;380(9841):572–80.

15. Do R, Willer CJ, Schmidt EM, Sengupta S, Gao C, Peloso GM, et al. Common variants associated with plasma triglycerides and risk for coronary artery disease. Nat Genet. 2013;45(11):1345–52.

16. Bulik-Sullivan B, Finucane HK, Anttila V, Gusev A, Day FR, Loh PR, et al. An atlas of genetic correlations across human diseases and traits. Nat Genet. 2015;47(11):1236–41.

17. Ellinghaus D, Jostins L, Spain SL, Cortes A, Bethune J, Han B, et al. Analysis of five chronic inflammatory diseases identifies 27 new associations and highlights disease-specific patterns at shared loci. Nat Genet. 2016;48(5):510–8.

18. Brainstorm Consortium, Anttila V, Bulik-Sullivan B, Finucane HK, Walters RK, Bras J, et al. Analysis of shared heritability in common disorders of the brain. Science. 2018;360(6395)

19. Chesmore K, Bartlett J, Williams SM. The ubiquity of pleiotropy in human disease. Hum Genet. 2018;137(1):39–44.

20. Verbanck M, Chen CY, Neale B, Do R. Detection of widespread horizontal pleiotropy in causal relationships inferred from Mendelian randomization between complex traits and diseases. Nat Genet. 2018;50(5):693–8.

21. Jordan DM, Verbanck M, Do R. The landscape of pervasive horizontal pleiotropy in human genetic variation is driven by extreme polygenicity of human traits and diseases. bioRxiv. 2018;doi: https://doi.org/10.1101/416545

22. Gandal MJ, Haney JR, Parikshak NN, Leppa V, Ramaswami G, Hartl C, et al. Shared molecular neuropathology across major psychiatric disorders parallels polygenic overlap. Science. 2018;359(6376):693–7.

23. Stringer S, Kahn RS, de Witte LD, Ophoff RA, Derks EM. Genetic liability for schizophrenia predicts risk of immune disorders. Schizophr Res. 2014;159:347–52.

24. Wang Q, Yang C, Gelernter J, Zhao H. Pervasive pleiotropy between psychiatric disorders and immune disorders revealed by integrative analysis of multiple GWAS. Hum Genet. 2015;134(11-12):1195–209.

25. Tylee DS, Sun J, Hess JL, Tahir MA, Sharma E, Malik R, et al. Genetic correlations among psychiatric and immune-related phenotypes based on genome-wide association data. Am J Med Genet B Neuropsychiatr Genet. 2018;177(7):641–57.

26. Duncan LE, Shen H, Ballon JS, Hardy KV, Noordsy DL, Levinson DF. Genetic correlation profile of schizophrenia mirrors epidemiological results and suggests link between polygenic and rare variant (22q11.2) cases of schizophrenia. Schizophr Bull. 2018;44(6):1350–61.

27. Pickrell JK, Berisa T, Liu JZ, Segurel L, Tung JY, Hinds DA. Detection and interpretation of shared genetic influences on 42 human traits. Nat Genet. 2016;48(7):709–17.

28. Andreassen OA, Harbo HF, Wang Y, Thompson WK, Schork AJ, Mattingsdal M, et al. Genetic pleiotropy between multiple sclerosis and schizophrenia but not bipolar disorder: differential involvement of immune-related gene loci. Mol Psychiatry. 2015;20:207–14.

29. Yin X, Wineinger NE, Wang K, Yue W, Norgren N, Wang L, et al. Common susceptibility variants are shared between schizophrenia and psoriasis in the Han Chinese population. J Psychiatry Neurosci. 2016;41(6):413–21.

30. Purcell SM, Wray NR, Stone JL, Visscher PM, O’Donovan MC, Sullivan PF, et al. Common polygenic variation contributes to risk of schizophrenia and bipolar disorder. Nature. 2009;460(7256):748–52.

31. Lee SH, Byrne EM, Hultman CM, Kahler A, Vinkhuyzen AA, Ripke S, et al. New data and an old puzzle: the negative association between schizophrenia and rheumatoid arthritis. Int J Epidemiol. 2015;44(5):1706–21.

32. Wray NR, Goddard ME, Visscher PM. Prediction of individual genetic risk to disease from genome-wide association studies. Genome Res. 2007;17(10):1520–8.

33. Fernando MM, Stevens CR, Walsh EC, De Jager PL, Goyette P, Plenge RM, et al. Defining the role of the MHC in autoimmunity: a review and pooled analysis. PLoS Genet. 2008;4:e1000024.

34. Franke A, McGovern DP, Barrett JC, Wang K, Radford-Smith GL, Ahmad T, et al. Genome-wide meta-analysis increases to 71 the number of confirmed Crohn’s disease susceptibility loci. Nat Genet. 2010;42(12):1118–25.

35. Jostins L, Ripke S, Weersma RK, Duerr RH, McGovern DP, Hui KY, et al. Host-microbe interactions have shaped the genetic architecture of inflammatory bowel disease. Nature. 2012;491(7422):119–24.

36. Beecham AH, Patsopoulos NA, Xifara DK, Davis MF, Kemppinen A, Cotsapas C, et al. Analysis of immune-related loci identifies 48 new susceptibility variants for multiple sclerosis. Nat Genet. 2013;45(11):1353–60.

37. Dubois PC, Trynka G, Franke L, Hunt KA, Romanos J, Curtotti A, et al. Multiple common variants for celiac disease influencing immune gene expression. Nat Genet. 2010;42(4):295–302.

38. Yang J, Ferreira T, Morris AP, Medland SE, Genetic IOANTGIANTC, DIAbetes GRAM-aDIAGRAMC, et al. Conditional and joint multiple-SNP analysis of GWAS summary statistics identifies additional variants influencing complex traits. Nat Genet. 2012;44(4):369–75, S1.

39. Zhu Z, Zhang F, Hu H, Bakshi A, Robinson MR, Powell JE, et al. Integration of summary data from GWAS and eQTL studies predicts complex trait gene targets. Nat Genet. 2016;48(5):481–7.

40. Wu Y, Zeng J, Zhang F, Zhu Z, Qi T, Zheng Z, et al. Integrative analysis of omics summary data reveals putative mechanisms underlying complex traits. Nat Commun. 2018;9(1):918.

41. 1000 Genomes Project Consortium, Auton A, Brooks LD, Durbin RM, Garrison EP, Kang HM, et al. A global reference for human genetic variation. Nature. 2015;526(7571):68–74.

42. Stahl EA, Wegmann D, Trynka G, Gutierrez-Achury J, Do R, Voight BF, et al. Bayesian inference analyses of the polygenic architecture of rheumatoid arthritis. Nat Genet. 2012;44(5):483–9.

43. Wood AR, Esko T, Yang J, Vedantam S, Pers TH, Gustafsson S, et al. Defining the role of common variation in the genomic and biological architecture of adult human height. Nat Genet. 2014;46(11):1173–86.

44. Lee SH, Goddard ME, Wray NR, Visscher PM. A better coefficient of determination for genetic profile analysis. Genet Epidemiol. 2012;36(3):214–24.

45. Finucane HK, Bulik-Sullivan B, Gusev A, Trynka G, Reshef Y, Loh PR, et al. Partitioning heritability by functional annotation using genome-wide association summary statistics. Nat Genet. 2015;47(11):1228–35.

46. Fairweather D, Frisancho-Kiss S, Rose NR. Sex differences in autoimmune disease from a pathological perspective. Am J Pathol. 2008;173(3):600–9.

47. Jones PA. Functions of DNA methylation: islands, start sites, gene bodies and beyond. Nat Rev Genet. 2012;13(7):484–92.

48. Bell AC, Felsenfeld G. Methylation of a CTCF-dependent boundary controls imprinted expression of the Igf2 gene. Nature. 2000;405(6785):482–5.

49. Eden S, Constancia M, Hashimshony T, Dean W, Goldstein B, Johnson AC, et al. An upstream repressor element plays a role in Igf2 imprinting. EMBO J. 2001;20(13):3518–25.

50. Davis S, Gale NW, Aldrich TH, Maisonpierre PC, Lhotak V, Pawson T, et al. Ligands for EPH-related receptor tyrosine kinases that require membrane attachment or clustering for activity. Science. 1994;266(5186):816–9.

51. Hruska M, Dalva MB. Ephrin regulation of synapse formation, function and plasticity. Mol Cell Neurosci. 2012;50(1):35–44.

52. Dustin ML. Signaling at neuro/immune synapses. J Clin Invest. 2012;122(4):1149–55.

53. Basu R, Huse M. Mechanical communication at the immunological synapse. Trends Cell Biol. 2017;27(4):241–54.

54. Sharfe N, Freywald A, Toro A, Dadi H, Roifman C. Ephrin stimulation modulates T cell chemotaxis. Eur J Immunol. 2002;32(12):3745–55.

55. Lu Q, Sun EE, Klein RS, Flanagan JG. Ephrin-B reverse signaling is mediated by a novel PDZ-RGS protein and selectively inhibits G protein-coupled chemoattraction. Cell. 2001;105(1):69–79.

56. U.S. Food and Drug Administration, Centre for Drug Evaluation and Research. Sprycel (dasatinib) NDA 21986/22072 approval letter, 2006 June 28. Retrieved 2018 November 4, from https://www.accessdata.fda.gov/drugsatfda_docs/nda/2006/021986s000_SprycelAPPR_OV.pdf.

57. ClinicalTrials.gov [Internet]. Bethesda (MD): National Library of Medicine (US). 2016 March 23. Identifier: NCT02717156, Combination therapy with Pembrolizumab and sEphB4-HSA in previously treated urothelial carcinoma. Available from: https://clinicaltrials.gov/ct2/show/NCT02717156.

58. ClinicalTrials.gov [Internet]. Bethesda (MD): National Library of Medicine (US). 2016 June 15. Identifier NCT02799485, EphB4-HSA in treating patients with Kaposi sarcoma. Available from: https://clinicaltrials.gov/ct2/show/NCT02799485.

59. ClinicalTrials.gov [Internet]. Bethesda (MD): National Library of Medicine (US). 2017 May 10. Identifier NCT03146871, Recombinant EphB4-HSA fusion protein and Azacitidine or Decitabine in treating patients with relapsed or refractory myelodysplastic syndrome, chronic myelomonocytic leukemia, or acute myeloid leukemia previously treated with a hypomethylating agent. Available from: https://clinicaltrials.gov/ct2/show/NCT03146871.

60. ClinicalTrials.gov [Internet] C. Bethesda (MD): National Library of Medicine (US). 2017 February 10. Identifier: NCT03049618, Recombinant EphB4-HSA fusion protein and Pembrolizumab in treating patients with locally advanced or metastatic non-small cell lung cancer or locally recurrent or metastatic head and neck squamous cell cancer. Available from: https://clinicaltrials.gov/ct2/show/NCT03049618.

61. Canellas F, Lin L, Julia MR, Clemente A, Vives-Bauza C, Ollila HM, et al. Dual cases of type 1 narcolepsy with schizophrenia and other psychotic disorders. J Clin Sleep Med. 2014;10(9):1011–8.

62. Arad M, Weiner I. Disruption of latent inhibition induced by ovariectomy can be reversed by estradiol and clozapine as well as by co-administration of haloperidol with estradiol but not by haloperidol alone. Psychopharmacology (Berl). 2009;206(4):731–40.

63. Zhu ML, Bakhru P, Conley B, Nelson JS, Free M, Martin A, et al. Sex bias in CNS autoimmune disease mediated by androgen control of autoimmune regulator. Nat Commun. 2016;7:11350.

64. Markle JG, Frank DN, Mortin-Toth S, Robertson CE, Feazel LM, Rolle-Kampczyk U, et al. Sex differences in the gut microbiome drive hormone-dependent regulation of autoimmunity. Science. 2013;339(6123):1084–8.

65. Zammit S, Rasmussen F, Farahmand B, Gunnell D, Lewis G, Tynelius P, et al. Height and body mass index in young adulthood and risk of schizophrenia: a longitudinal study of 1 347 520 Swedish men. Acta Psychiatr Scand. 2007;116(5):378–85.

66. Takayanagi Y, Petersen L, Laursen TM, Cascella NG, Sawa A, Mortensen PB, et al. Risk of schizophrenia spectrum and affective disorders associated with small for gestational age birth and height in adulthood. Schizophr Res. 2014;160(1-3):230–2.

67. Boyle EA, Li YI, Pritchard JK. An expanded view of complex traits: from polygenic to omnigenic. Cell. 2017;169(7):1177–86.

68. Trynka G, Hunt KA, Bockett NA, Romanos J, Mistry V, Szperl A, et al. Dense genotyping identifies and localizes multiple common and rare variant association signals in celiac disease. Nat Genet. 2011;43(12):1193–201.

69. Liu JZ, van Sommeren S, Huang H, Ng SC, Alberts R, Takahashi A, et al. Association analyses identify 38 susceptibility loci for inflammatory bowel disease and highlight shared genetic risk across populations. Nat Genet. 2015;47(9):979–86.

70. Hinks A, Cobb J, Marion MC, Prahalad S, Sudman M, Bowes J, et al. Dense genotyping of immune-related disease regions identifies 14 new susceptibility loci for juvenile idiopathic arthritis. Nat Genet. 2013;45(6):664–9.

71. Faraco J, Lin L, Kornum BR, Kenny EE, Trynka G, Einen M, et al. ImmunoChip study implicates antigen presentation to T cells in narcolepsy. PLoS Genet. 2013;9(2):e1003270.

72. Stahl EA, Raychaudhuri S, Remmers EF, Xie G, Eyre S, Thomson BP, et al. Genome-wide association study meta-analysis identifies seven new rheumatoid arthritis risk loci. Nat Genet. 2010;42(6):508–14.

73. Bentham J, Morris DL, Cunninghame Graham DS, Pinder CL, Tombleson P, Behrens TW, et al. Genetic association analyses implicate aberrant regulation of innate and adaptive immunity genes in the pathogenesis of systemic lupus erythematosus. Nat Genet. 2015;47(12):1457–64.

74. Onengut-Gumuscu S, Chen WM, Burren O, Cooper NJ, Quinlan AR, Mychaleckyj JC, et al. Fine mapping of type 1 diabetes susceptibility loci and evidence for colocalization of causal variants with lymphoid gene enhancers. Nat Genet. 2015;47(4):381–6.

75. Cordell HJ, Han Y, Mells GF, Li Y, Hirschfield GM, Greene CS, et al. International genome-wide meta-analysis identifies new primary biliary cirrhosis risk loci and targetable pathogenic pathways. Nat Commun. 2015;6:8019.

76. Strange A, Capon F, Spencer CC, Knight J, Weale ME, Allen MH, et al. A genome-wide association study identifies new psoriasis susceptibility loci and an interaction between HLA-C and ERAP1. Nat Genet. 2010;42(11):985–90.

77. Radstake TR, Gorlova O, Rueda B, Martin JE, Alizadeh BZ, Palomino-Morales R, et al. Genome-wide association study of systemic sclerosis identifies CD247 as a new susceptibility locus. Nat Genet. 2010;42(5):426–9.

78. Jin Y, Andersen G, Yorgov D, Ferrara TM, Ben S, Brownson KM, et al. Genome-wide association studies of autoimmune vitiligo identify 23 new risk loci and highlight key pathways and regulatory variants. Nat Genet. 2016;48(11):1418–24.

79. Yang J, Lee SH, Goddard ME, Visscher PM. GCTA: a tool for genome-wide complex trait analysis. Am J Hum Genet. 2011;88(1):76–82.

80. Powell JE, Henders AK, McRae AF, Caracella A, Smith S, Wright MJ, et al. The Brisbane Systems Genetics Study: genetical genomics meets complex trait genetics. PLoS One. 2012;7(4):e35430.

81. Chen BH, Marioni RE, Colicino E, Peters MJ, Ward-Caviness CK, Tsai PC, et al. DNA methylation-based measures of biological age: meta-analysis predicting time to death. Aging (Albany NY). 2016;8(9):1844–65.

82. Lloyd-Jones LR, Holloway A, McRae A, Yang J, Small K, Zhao J, et al. The genetic architecture of gene expression in peripheral blood. Am J Hum Genet. 2017;100(2):371.

83. R Core Team. R: A language and environment for statistical computing. R Foundation for Statistical Computing. 2012;Vienna, Austria. ISBN 3-900051-07-0, URL http://www.R-project.org/.

84. Nakazawa M. fmsb: Functions for medical statistics book with some demographic data. R package version 051. 2014http://CRAN.R-project.org/package=fmsb.

85. Psychiatric GWAS Consortium Bipolar Disorder Working Group. Large-scale genome-wide association analysis of bipolar disorder identifies a new susceptibility locus near ODZ4. Nat Genet. 2011;43(10):977–83.

86. Bulik-Sullivan BK, Loh PR, Finucane HK, Ripke S, Yang J, Patterson N, et al. LD Score regression distinguishes confounding from polygenicity in genome-wide association studies. Nat Genet. 2015;47(3):291–5.

87. Abecasis GR, Auton A, Brooks LD, DePristo MA, Durbin RM, Handsaker RE, et al. An integrated map of genetic variation from 1,092 human genomes. Nature. 2012;491(7422):56–65.

88. International HapMap 3 Consortium, Altshuler DM, Gibbs RA, Peltonen L, Altshuler DM, Gibbs RA, et al. Integrating common and rare genetic variation in diverse human populations. Nature. 2010;467(7311):52–8.

89. Purcell S, Cherny SS, Sham PC. Genetic Power Calculator: design of linkage and association genetic mapping studies of complex traits. Bioinformatics. 2003;19(1):149–50.

90. Palla L, Dudbridge F. A fast method that uses polygenic scores to estimate the variance explained by genome-wide marker panels and the proportion of variants affecting a trait. Am J Hum Genet. 2015;97(2):250–9.

91. Dudbridge F. Power and predictive accuracy of polygenic risk scores. PLoS Genet. 2013;9(3):e1003348.

92. Betz RC, Petukhova L, Ripke S, Huang H, Menelaou A, Redler S, et al. Genome-wide meta-analysis in alopecia areata resolves HLA associations and reveals two new susceptibility loci. Nat Commun. 2015;6:5966.

93. Cortes A, Hadler J, Pointon JP, Robinson PC, Karaderi T, Leo P, et al. Identification of multiple risk variants for ankylosing spondylitis through high-density genotyping of immune-related loci. Nat Genet. 2013;45(7):730–8.

94. Chu X, Pan CM, Zhao SX, Liang J, Gao GQ, Zhang XM, et al. A genome-wide association study identifies two new risk loci for Graves’ disease. Nat Genet. 2011;43(9):897–901.

95. Gutierrez-Achury J, Zhernakova A, Pulit SL, Trynka G, Hunt KA, Romanos J, et al. Fine mapping in the MHC region accounts for 18% additional genetic risk for celiac disease. Nat Genet. 2015;47(6):577–8.

96. Goyette P, Boucher G, Mallon D, Ellinghaus E, Jostins L, Huang H, et al. High-density mapping of the MHC identifies a shared role for HLA-DRB1*01:03 in inflammatory bowel diseases and heterozygous advantage in ulcerative colitis. Nat Genet. 2015;47(2):172–9.

97. Patsopoulos NA, Barcellos LF, Hintzen RQ, Schaefer C, van Duijn CM, Noble JA, et al. Fine-mapping the genetic association of the major histocompatibility complex in multiple sclerosis: HLA and non-HLA effects. PLoS Genet. 2013;9(11):e1003926.

98. Tafti M, Hor H, Dauvilliers Y, Lammers GJ, Overeem S, Mayer G, et al. DQB1 locus alone explains most of the risk and protection in narcolepsy with cataplexy in Europe. Sleep. 2014;37(1):19–25.

99. Liu JZ, Almarri MA, Gaffney DJ, Mells GF, Jostins L, Cordell HJ, et al. Dense fine-mapping study identifies new susceptibility loci for primary biliary cirrhosis. Nat Genet. 2012;44(10):1137–41.

100. Liu JZ, Hov JR, Folseraas T, Ellinghaus E, Rushbrook SM, Doncheva NT, et al. Dense genotyping of immune-related disease regions identifies nine new risk loci for primary sclerosing cholangitis. Nat Genet. 2013;45(6):670–5.

101. Okada Y, Han B, Tsoi LC, Stuart PE, Ellinghaus E, Tejasvi T, et al. Fine mapping major histocompatibility complex associations in psoriasis and its clinical subtypes. Am J Hum Genet. 2014;95(2):162–72.

102. Raychaudhuri S, Sandor C, Stahl EA, Freudenberg J, Lee HS, Jia X, et al. Five amino acids in three HLA proteins explain most of the association between MHC and seropositive rheumatoid arthritis. Nat Genet. 2012;44(3):291–6.

103. Lessard CJ, Li H, Adrianto I, Ice JA, Rasmussen A, Grundahl KM, et al. Variants at multiple loci implicated in both innate and adaptive immune responses are associated with Sjogren’s syndrome. Nat Genet. 2013;45(11):1284–92.

104. Kim K, Bang SY, Lee HS, Okada Y, Han B, Saw WY, et al. The HLA-DRbeta1 amino acid positions 11-13-26 explain the majority of SLE-MHC associations. Nat Commun. 2014;5:5902.

105. Mayes MD, Bossini-Castillo L, Gorlova O, Martin JE, Zhou X, Chen WV, et al. Immunochip analysis identifies multiple susceptibility loci for systemic sclerosis. Am J Hum Genet. 2014;94(1):47–61.

106. Hu X, Deutsch AJ, Lenz TL, Onengut-Gumuscu S, Han B, Chen WM, et al. Additive and interaction effects at three amino acid positions in HLA-DQ and HLA-DR molecules drive type 1 diabetes risk. Nat Genet. 2015;47(8):898–905.

