## Supplementary material for "Cross-disorder analysis of schizophrenia and 19 immune diseases reveals genetic correlation": Pouget2019_SupplementaryInformation

**Supplementary Information**

| **Supplementary Methods** | | Page 2 |
| --- | --- | --- |
| **Supplementary Tables** | | |
| **S1 Table** | Previous investigations of genetic overlap between schizophrenia and immune-mediated disease | Page 4 |
| **S2 Table** | Pleiotropic associations between DNAm sites, transcripts and schizophrenia from SMR and HEIDI analyses | Page 7 |
| **S3 Table** | Association of polygenic risk scores excluding HLA variants for height (negative control) and 14 autoimmune diseases with schizophrenia case-status | Page 7 |
| **S4 Table** | Association of polygenic risk scores including top HLA variant for 14 autoimmune diseases with schizophrenia case status | Page 7 |
| **S5 Table** | Sex-stratified association of non-HLA polygenic risk scores for 14 autoimmune diseases with schizophrenia case status | Page 7 |
| **S6 Table** | Parameters used for polygenic risk scoring statistical power calculations | Page 8 |
| **Supplementary Figures** | | |
| **S1 Fig** | Effect of immune disease risk variants on schizophrenia susceptibility | Page 9 |
| **S2 Fig** | Association of immune risk SNPs in schizophrenia | Page 12 |
| **S3 Fig** | Identification of pleiotropic HLA variants | Page 13 |
| **S4 Fig** | Colocalization of GWAS association signals for schizophrenia and immune diseases | Page 13 |
| **S5 Fig** | Statistical power to detect genetic overlap between schizophrenia and immune-mediated diseases using polygenic risk scoring | Page 17 |
| **S6 Fig** | Prediction of schizophrenia using genetic liability for 14 immune-mediated diseases excluding HLA variants | Page 18 |
| **References** |  | Page 20 |
| **Members of the Schizophrenia Working Group of the   Psychiatric Genomics Consortium** | | Page 24 |

**Supplementary Methods**

**Defining immune disease risk loci using ImmunoBase curations**

For non-HLA risk loci, curated GWAS results were obtained from ImmunoBase (TAB files from http://www.immunobase.org/downloads/regions-files-archives/2015-06-07; accessed 7 June 2015). For each of the 19 immune diseases with available curations, we pruned the list of index SNPs using the 1000 Genomes Phase 1 European reference panel for LD to list all pairs of SNPs with r^2^ > 0.1 using the PLINK [1] command --r2 --ld-window-r2 0.1, and kept the most strongly associated SNP. To ensure complete independence we also removed SNPs that were annotated in the same chromosomal region (e.g. 1p13.2) by ImmunoBase, again keeping the most strongly associated index SNP.

**Collective association of risk variants for autoimmune disease in schizophrenia**

To evaluate the collective association of SNPs associated with autoimmune disease with schizophrenia, we pruned the list of 563 unique SNPs across autoimmune disorders (pairwise r^2^ < 0.1 based on 1000 Genomes Phase I European reference panel) using PLINK [1] with the --r2 --ld-window-r2 0.1 function. In total, we analyzed 261 immune risk SNPs for which LD Score and MAF information was available in the European LD Score database [2]. Next, we evaluated the p-values of these LD-independent immune risk SNPs in the schizophrenia dataset. We quantified enrichment of autoimmune SNP associations in schizophrenia using the genomic inflation value λ. We obtained an empirical enrichment p-value by comparing λ_observed_ to λ values obtained in 1,000 permutations using the following procedure: we randomly selected 1,000 sets of SNPs that matched the immune SNP set for minor allele frequency (MAF, using 10 bins at increments of 0.05: i.e. 0<MAF≤0.05, 0.05<MAF≤0.10, 0.10<MAF≤0.15,…) and linkage disequilibrium (LD) score (using 10 bins: 0<LD score≤25, 25<LD score≤40, 40<LD score≤55, 55<LD score≤70, 70<LD score≤85, 85<LD score≤100, 100<LD score≤125, 125<LD score≤150, 150<LD score≤200, LD score>200) [3]. MAFs and LD scores were obtained from the LD Score database of Bulik-Sullivan *et al.* (see **URLs**), which uses the 1000 Genomes Phase I European population as a reference.

**Polygenic risk scoring**

Each of the training (immune GWAS) datasets underwent the following quality control (QC) steps: SNPs on non-autosomal chromosomes (X, Y, M) were removed, SNPs with MAF < 0.01 were removed if MAF was available in the training dataset, SNPs with INFO < 0.90 were removed if INFO was available in the training dataset, SNPs with missing p-value or OR were removed, symmetrical SNPs were removed. For training datasets where the effect size was provided as an OR, this was converted to ß coefficients (log_10_(OR)). Risk alleles were recoded such that all risk alleles corresponded to increased risk of diseases (all ß > 0). After QC, each of the training dataset (immune GWAS) SNP sets was merged with the target schizophrenia GWAS SNP set. Clumping was performed using PLINK [1], with the 1000 Genomes Phase I European population as a reference, to retain SNPs with r^2^ < 0.1 within 1,000 kb windows, while filtering for the highest significance levels within LD blocks (using options --clump-p1 1 --clump-p2 1 --clump-r2 0.1 --clump-kb 1000).

Using the clumped SNP list for each of the training datasets, we calculated PRSs in PLINK [1] with the --score function. We standardized the PRSs to have a mean of 0 and s.d. of 1 within each schizophrenia cohort to allow comparison of all PRSs on the same scale, as previously described [4, 5]. We excluded eight cohorts (caws, clm2, clo3, cou3, irwt, lacw, pewb, pews; see **Supplementary Table 1** of initial GWAS for full sample details [6]) that included controls from the Wellcome Trust Case Control Consortium (WTCCC), due to the use of at least a subset of these samples in the training datasets. We then performed logistic regression on the total sample in R [7] using the stats package [7] to evaluate the association between schizophrenia case status and PRSs for each immune disease. The variance in schizophrenia case status explained by the PRSs was estimated using the deviation in liability-scale R^2^ between a null model (which included 10 ancestry-informative principal components and study site) to the full model (which included PRSs in addition to these covariates). Liability-scale R^2^ was calculated as previously described [8], assuming a population prevalence of schizophrenia of 1%. Liability-scale R^2^ was used as the primary effect size measure, because it has been described as a better coefficient of determination in PRS studies because it is consistent with the underlying scale, independent of sample parameters (such as disease prevalence and proportion of cases), and easily interpretable in relation to true heritability [8]. We also calculated Nagelkerke’s pseudo-R^2^, in R [7] using the fmsb package [9]. Similar to previous studies, statistical significance of the PRSs was estimated based on their logistic regression coefficient [10, 11]. We used Bonferroni correction to establish a significance threshold of 0.05/19=2.6x10^-3^ (to correct for 19 immune-mediated diseases tested for genetic overlap with schizophrenia), and considered immune diseases with PRSs significant at this threshold to show significant genetic overlap with schizophrenia.

S1 Table. Previous investigations of genetic overlap between schizophrenia and immune-mediated disease

|  | **Study** | **GWAS Samples** | **Method** | **Result**^a^ |
| --- | --- | --- | --- | --- |
| CEL | Tylee 2018 [12] | Schizophrenia PGC2 [6]  CEL [13] | LDSC | n.s. |
| CRO | Purcell 2009 [10] | Schizophrenia ISC [10] ***training***  CRO [14] ***target*** | PRS | n.s. |
|  | Cross-Disorder Group  of the PGC 2013 [15] | Schizophrenia PGC1 - shared samples [15]  CRO - shared samples [16] | REML | n.s. |
|  | Stringer 2014 [5] | Schizophrenia PGC1 - caws [6] ***training***  CRO [14] ***target*** | PRS | +* |
|  | Bulik-Sullivan 2015 [2] | Schizophrenia PGC2 - Asian [6]  CRO [17] | LDSC | n.s. |
|  | Wang 2015 [18] | Schizophrenia PGC Cross-Disorder [15]  CRO [16] | GPA | +* |
|  | Pickrell 2016 [19] | Schizophrenia PGC2 [6]  CRO [20] | GWAS-PW | + |
|  | Tylee 2018 [12] | Schizophrenia PGC2 [6]  CRO [16] | LDSC | +* |
|  | Duncan 2018 [21] | Schizophrenia  CRO [17] | LDSC | + |
| MS | Andreassen 2014 [22] | MS [23] ***training***  Schizophrenia PGC1 [24] ***target*** | CFDR | +* |
|  | Wang 2015 [18] | Schizophrenia PGC Cross-Disorder [11]  MS [25] | GPA | n.s. |
|  | Tylee 2018 [12] | Schizophrenia PGC2 [6]  MS [23] | LDSC | n.s. |
| PBC | Tylee 2018 [12] | Schizophrenia PGC2 [6]  PBC [26] | LDSC | +* |
| PSO | Wang 2015 [18] | Schizophrenia PGC Cross-Disorder [11]  PSO [27] | GPA | n.s. |
|  | Yin 2016 [28] | SCZ [29]  PSO [30] | REML | +* |
|  | Tylee 2018 [12] | Schizophrenia PGC2 [6]  PSO [31] | LDSC | + |
| RA | Purcell 2009 [10] | Schizophrenia ISC [10] ***training***  RA [14] ***target*** | PRS | n.s. |
|  | Stringer 2014 [5] | Schizophrenia PGC1 - caws [6] ***training***  RA [14] ***target*** | PRS | +* |
|  | Lee 2015 [32] | Schizophrenia PGC1+Swe [24, 33]  RA [34, 35] | REML | - * |
|  | Euesden 2015 [36] | Schizophrenia PGC1.5 [33]  RA [14] | PRS | n.s. |
|  | Bulik-Sullivan 2015 [2] | Schizophrenia PGC2 - Asian [6]  RA [34] | LDSC | n.s. |
|  | Wang 2015 [18] | Schizophrenia PGC Cross-Disorder [11]  RA [34] | GPA | + |
|  | Pickrell 2016 [19] | Schizophrenia PGC2 [6]  RA [35] | GWAS-PW | n.s. |
|  | Tylee 2018 [12] | Schizophrenia PGC2 [6]  RA [35] | LDSC | n.s. |
|  | Duncan 2018 [21] | Schizophrenia PGC2 [6]  RA [35] | LDSC | n.s. |
| SLE | Wang 2015 [18] | Schizophrenia PGC Cross-Disorder [11]  SLE [37] | GPA | + |
|  | Tylee 2018 [12] | Schizophrenia PGC2 [6]  SLE [38] | LDSC | +* |
| SSC | Tylee 2018 [12] | Schizophrenia PGC2 [6]  SSC [39] | LDSC | n.s. |
| T1D | Purcell 2009 [10] | Schizophrenia ISC [10] ***training***  T1D [14] **target** | PRS | n.s. |
|  | Stringer 2014 [5] | Schizophrenia PGC1 - caws [6] ***training***  T1D [14] ***target*** | PRS | +* |
|  | Wang 2015 [18] | Schizophrenia PGC Cross-Disorder [11]  T1D [40] | GPA | n.s. |
|  | Tylee 2018 [12] | Schizophrenia PGC2 [6]  T1D [41] | LDSC | n.s. |
| UC | Bulik-Sullivan 2015 [2] | Schizophrenia PGC2 - Asian [6]  UC [17] | LDSC | n.s. |
|  | Wang 2015 [18] | Schizophrenia PGC Cross-Disorder [11]  UC [42] | GPA | +* |
|  | Pickrell 2016 [19] | Schizophrenia PGC2 [6]  UC [20] | GWAS-PW | +* |
|  | Tylee 2018 [12] | Schizophrenia PGC2 [6]  UC [42] | LDSC | +* |
|  | Duncan 2018 [21] | Schizophrenia PGC2 [6]  UC [17] | LDSC | + |

^a^Only those results that were robust to exclusion of the major histocompatibility complex (MHC) region are presented as significant in this table. n.s., not significant; +, positive genetic correlation between schizophrenia and the immune disease of interest; -, negative genetic correlation between schizophrenia and the immune disease of interest; *, finding survives multiple testing correction; CEL, celiac disease; CFDR, conditional false discovery rate [43]; CRO, Crohn’s disease; GPA, Genetic analysis incorporating pleiotropy and annotation [44]; GWAS-PW, Bayesian framework for estimation of the proportion of genetic variants influencing two traits [19]; ISC, International Schizophrenia Consortium; LDSC, cross-trait LD Score regression [2]; MS, multiple sclerosis; PRS, polygenic risk scoring [45]; PSO, psoriasis, RA, rheumatoid arthritis; REML, restricted maximum likelihood [46]; SLE, systemic lupus erythematosus; T1D, type 1 diabetes; UC, ulcerative colitis.

S2 Table. Pleiotropic associations between DNAm sites, transcripts and schizophrenia from SMR and HEIDI analyses

See attached Excel file

S3 Table. Association of polygenic risk scores excluding HLA variants for height (negative control) and 14 autoimmune diseases with schizophrenia case-status

See attached Excel file

S4 Table. Association of polygenic risk scores including top HLA variant for 14 autoimmune diseases with schizophrenia case-status

See attached Excel file

S5 Table. Sex-stratified association of non-HLA polygenic risk scores for 14 autoimmune diseases with schizophrenia case-status

See attached Excel file

S6 Table. Parameters used for polygenic risk scoring power calculations

| **Disease** | **Prevalence** | **N Total** | **Case/Control  Sampling Proportion** | ***v*_g_^a^** | **N SNPs genotyped in both training and target** |
| --- | --- | --- | --- | --- | --- |
| SCZ | 0.01 [36] | 37,655 | 0.45 |  |  |
| CEL | 0.01 [47] | 24,269 [48] | 0.50 [48] | 0.32 [49] | 19,698 |
| CRO | 0.005 [50] | 20,883 [17] | 0.42 [17] | 0.26 [50] | 114,950 |
| IBD | 0.0075 [50] | 34,652 [17] | 0.59 [17] | 0.20 | 116,346 |
| JIA | 0.00025 [51] | 9,302^b^ [52] | 0.08 [52] | 0.18 [52] | 20,337 |
| MS | 0.001 [53] | 38,589 [54] | 0.38 [54] | 0.30 [55] | 21,818 |
| NAR | 0.0002 [56] | 12,307 [56] | 0.15 [56] | 0.20 | 19,866 |
| PBC | 0.0004 [57] | 11,375 [26] | 0.25 [26] | 0.20 | 97,806 |
| PSO | 0.02 [58] | 7,353 [59] | 0.30 [59] | 0.35^c^ [60] | 107,002 |
| RA | 0.01 [49] | 25,708 [34] | 0.22 [34] | 0.33 [49] | 126,049 |
| SLE | 0.00031^d^ [37] | 10,995 [38] | 0.37 [38] | 0.20 | 264,374 |
| SSC | 0.0001 [61] | 4,963^e^ [39] | 0.30 [39] | 0.09 [61] | 66,402 |
| T1D | 0.004 [62] | 22,175 [63] | 0.42 [63] | 0.20 | 20,835 |
| UC | 0.0025 [50] | 27,432 [17] | 0.34 [17] | 0.19 [50] | 120,720 |
| VIT | 0.004 [64] | 15,899 [65] | 0.09 [65] | 0.20 | 257,654 |

Where available, prevalence estimates used in the autoimmune GWASs were used. Otherwise, alternative reference is cited; ^a^Obtained from literature where possible. Otherwise, a variance of 0.20 was assumed based on average variances explained across many complex traits [66], [67]; ^b^Only the UK cohort from this study was available for analysis; ^c^Proportion of trait variance explained by genetic effects estimated in Chinese population; ^d^Prevalence estimate for European females; ^e^Only the US cohort from this study was available for analysis. CEL, celiac disease; CRO, Crohn’s disease; IBD, inflammatory bowel disease; JIA, juvenile idiopathic arthritis; MS, multiple sclerosis; NAR, narcolepsy; PBC, primary biliary cirrhosis; PSO, psoriasis; RA, rheumatoid arthritis; SCZ, schizophrenia; SLE, systemic lupus erythematosus; SSC, systemic sclerosis; T1D, type 1 diabetes; UC, ulcerative colitis; VIT, vitiligo; v_g_, proportion of trait variance explained by genetic effects in the training sample.

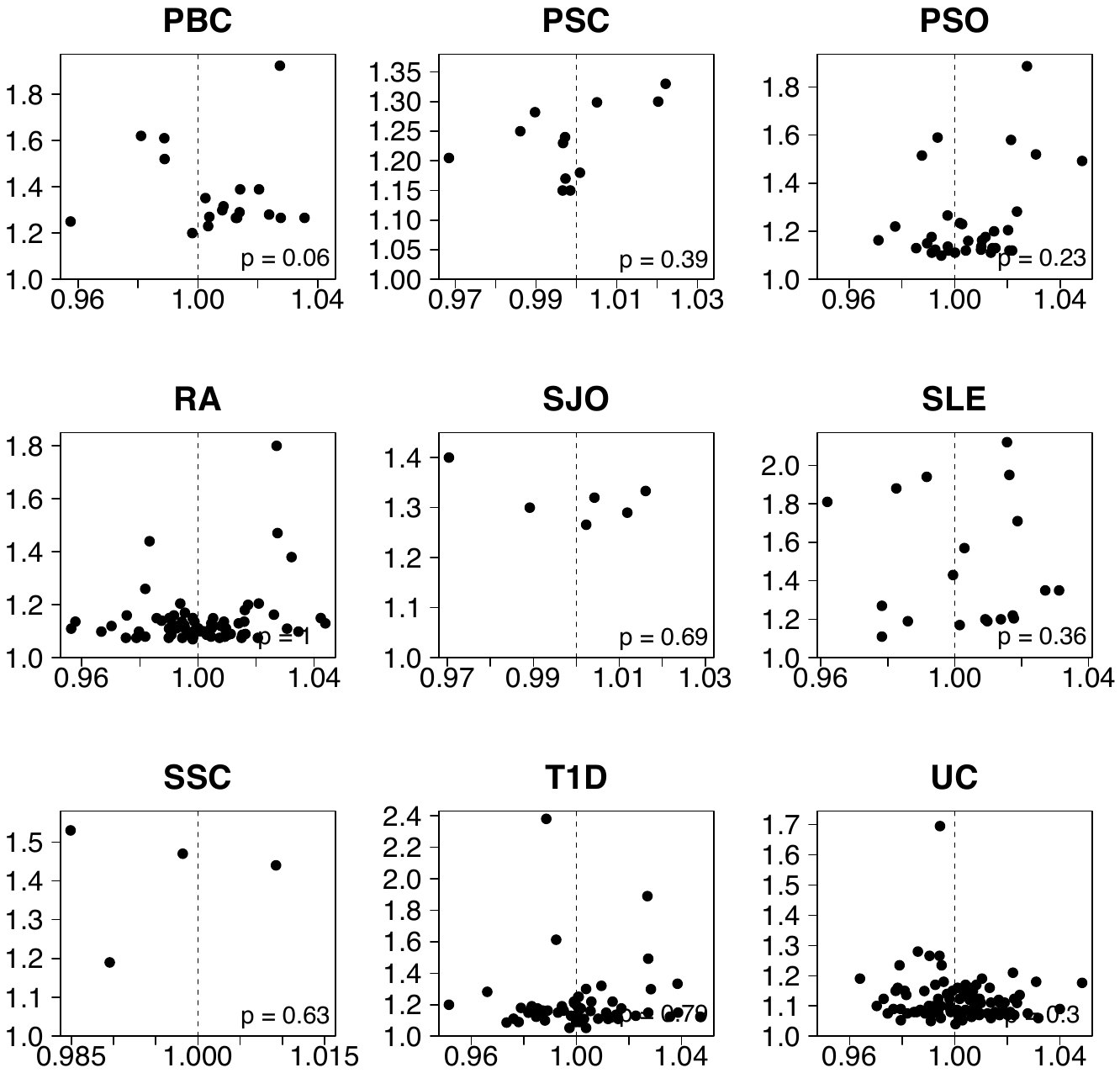

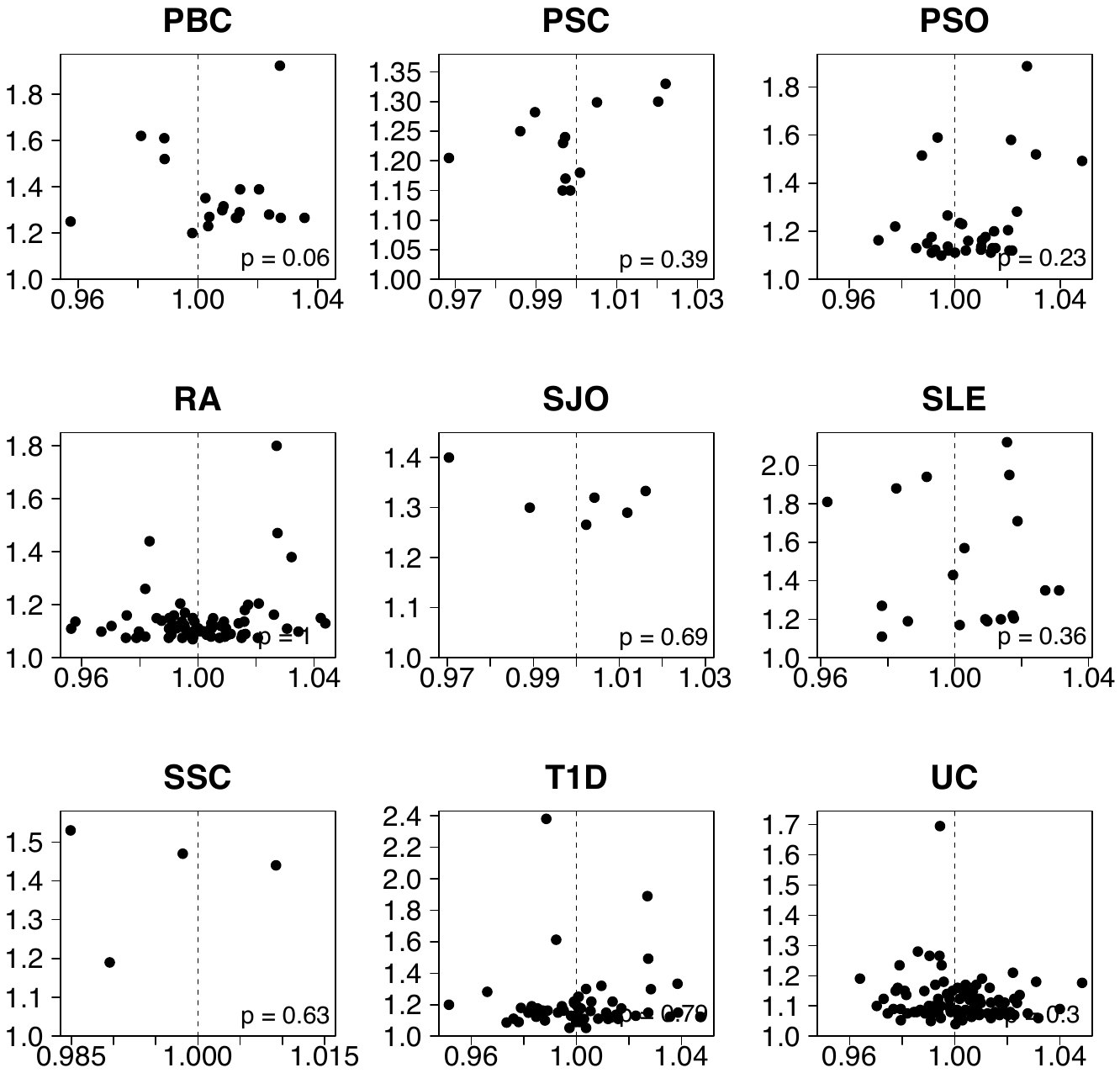

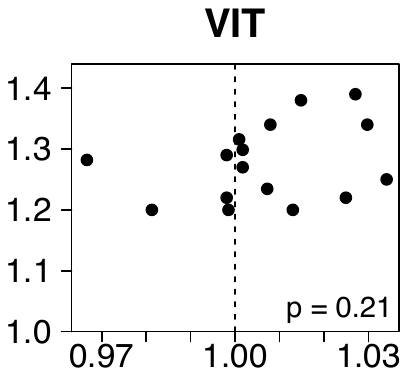

S1 Fig. Effect of immune disease risk variants on schizophrenia susceptibility

Genome-wide significant risk SNPs outside of the HLA region for 19 immune-mediated diseases were obtained from Immunobase, and evaluated for their association with schizophrenia. x-axis corresponds to the effect size (odds ratio, OR) of a SNP in schizophrenia, y-axis corresponds to the effect size (OR) in the immune disease of interest. P-values correspond to shared direction of effect for risk alleles between each of the immune diseases and schizophrenia, evaluated using the binomial sign test. AA, alopecia areata; AS, ankylosing spondylitis; ATD, autoimmune thyroid disease; CEL, celiac disease; CRO, Crohn’s disease; IBD, inflammatory bowel disease; JIA, juvenile idiopathic arthritis; MS, multiple sclerosis; NAR, narcolepsy; PBC, primary biliary cirrhosis; PSC, primary sclerosing cholangitis; PSO, psoriasis, RA, rheumatoid arthritis; SJO, Sjögren’s syndrome, SLE, systemic lupus erythematosus; SSC, systemic sclerosis; T1D, type 1 diabetes; UC, ulcerative colitis; VIT, vitiligo.

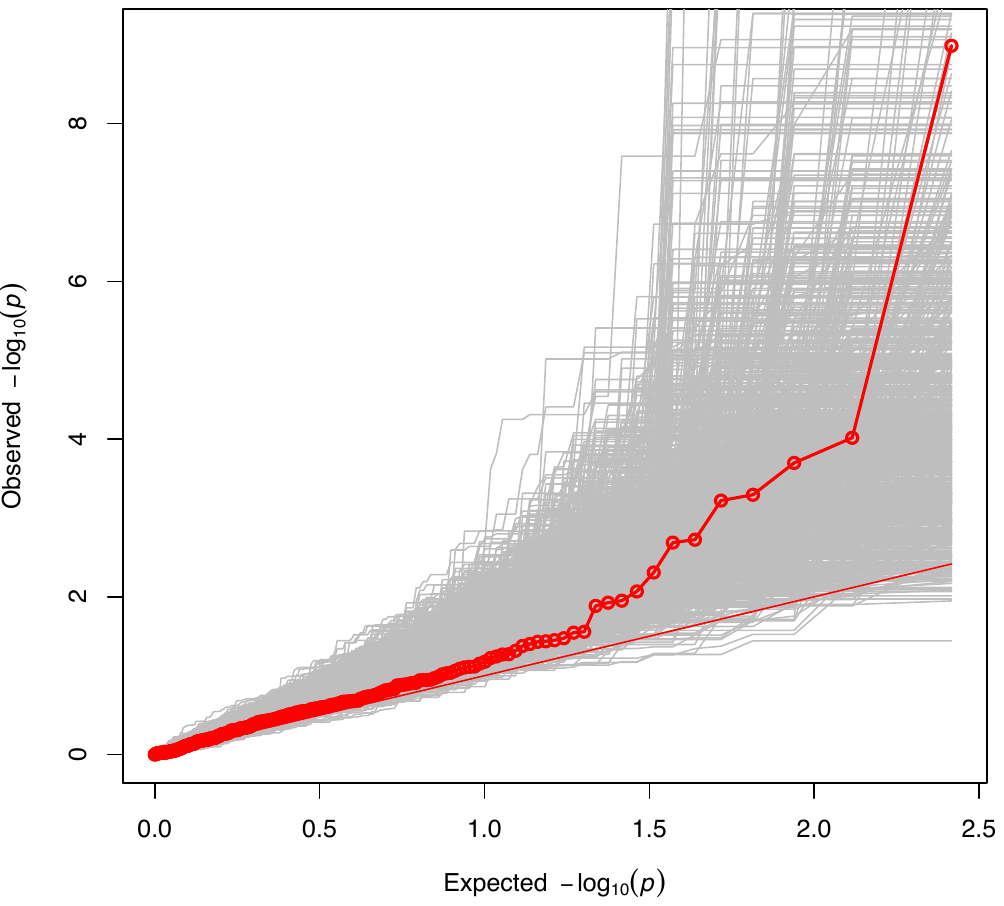

S2 Fig. Association of immune risk SNPs in schizophrenia

261 LD-independent non-HLA variants associated with an immune-mediated disease at genome-wide significance were evaluated for association with schizophrenia using publicly available summary statistics from the largest published schizophrenia GWAS [6]. Red points indicate the observed –log_10_(*p*) association of the 261 immune risk SNPs in schizophrenia. Gray lines indicate the observed –log_10_(*p*) association of 1,000 randomly selected sets of 261 SNPs, matched to the immune risk SNP set based on MAF and LD Score. An empirical p-value for *en masse* enrichment of the immune risk SNPs was obtained by comparing the number of permuted SNP sets with a larger λ than the observed λ of 1.46 for the 261 immune risk SNPs with available LD Score and MAF information (p=0.66)

**S3 Fig. Identification of pleiotropic HLA variants**

The most strongly associated HLA variant for each of the 19 immune-mediated diseases was evaluated for association with schizophrenia, using summary statistics from HLA imputation and association testing in the schizophrenia dataset as previously described (9). x-axis corresponds to position in the classical HLA region, shown in megabase pairs (Mb); y-axis corresponds to the HLA variant’s odds ratio (OR) in schizophrenia. Dashed line indicates OR=1. Size of the circle corresponds to the HLA variant’s OR in the immune disease of interest, which is indicated in the circle center. Red circles indicate HLA variants associated with schizophrenia above the Bonferroni significance threshold (p<8.6x10-5). Gene map below indicates location of coding HLA genes (coloured lines); gray lines correspond to non-HLA genes in the region. Disease abbreviations as defined in **Table 1**.

**S4 Fig. Colocalization of GWAS association signals for schizophrenia and immune diseases**

Regional association plots for schizophrenia [6] constructed using LocusZoom [68] at five potential pleiotropic loci: rs296547 (a), rs6738825 (b), rs13126505 (c), rs1734907 (d), and rs13126505 (e). These SNPs of interest are curated in ImmunoBase as risk variants for celiac disease (rs296547), Crohn’s disease (rs6738825, rs13126505, rs1734907), and multiple sclerosis (rs7132277) based on genome-wide association with these traits in previous studies. In the setting of a true pleiotropic role for these SNPs in schizophrenia and immune disease, no significant associations with schizophrenia should remain after conditioning on the SNP of interest (statistically, all p>8.6x10^-5^). Upper plot shows baseline association with schizophrenia, lower plot shows association with schizophrenia after conditioning on the SNP of interest using conditional and joint analysis (COJO) as implemented in GCTA [69]. Colour indicates LD with the SNP of interest, using 1000 Genomes Phase 3 CEU population as a reference panel [70].

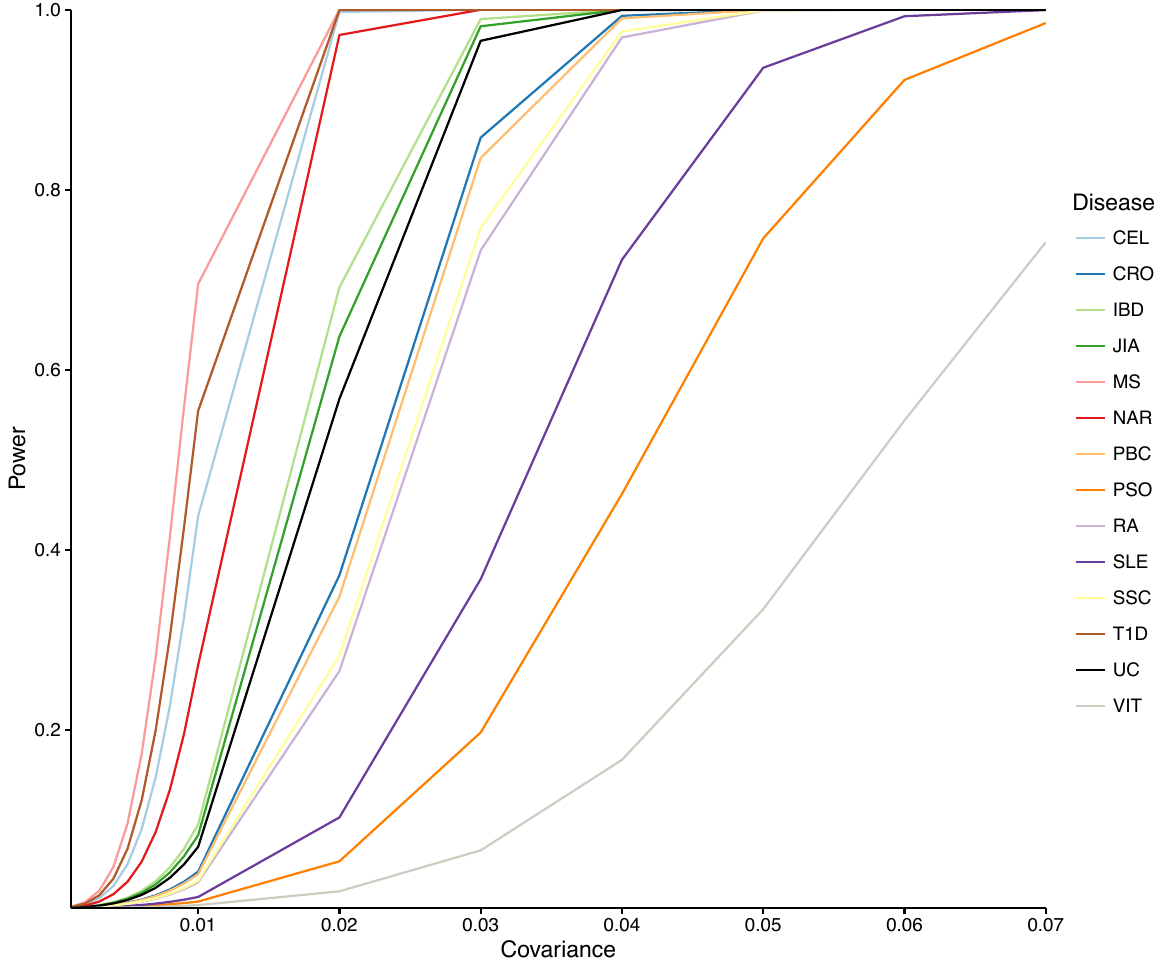

S5 Fig. Statistical power to detect genetic overlap between schizophrenia and immune-mediated diseases using polygenic risk scoring

Power was calculated as a function of the sizes of the training and target samples, disease prevalence, and explained genetic variance using a previously described quantitative genetics model [71] (see **S5 Table** for parameters). Power was calculated assuming a significance threshold of p_T_<1 for inclusion of SNPs in the PRS (i.e., including all SNPs). The significance threshold was adjusted using the Bonferroni correction accounting for 28 tests (14 diseases tested in both sexes). Disease abbreviations as defined in **S5 Table**.

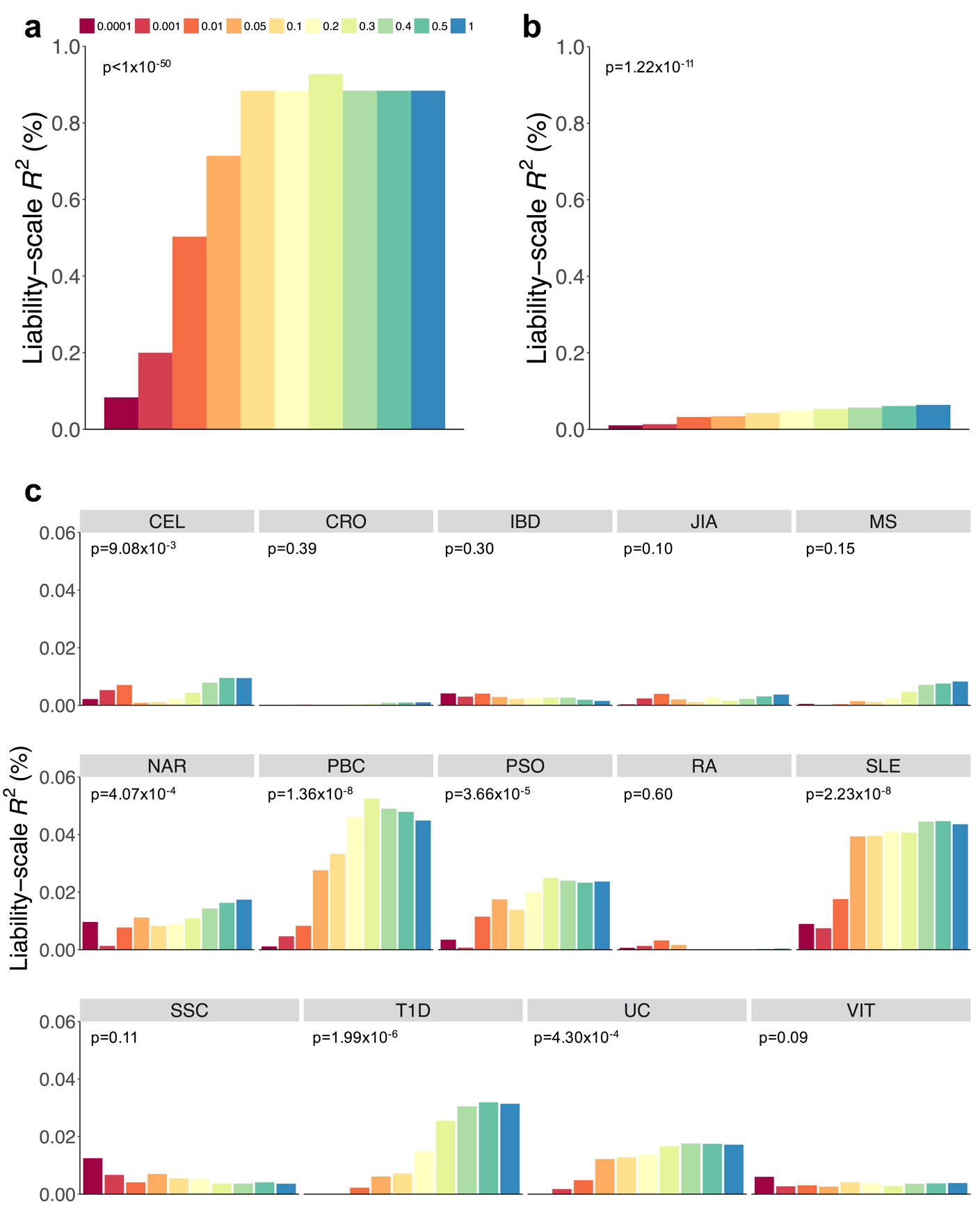

S6 Fig. Prediction of schizophrenia using genetic liability for 14 immune-mediated diseases excluding HLA variants

Prediction of schizophrenia case-status using polygenic risk scores constructed for bipolar disorder, a positive control (a), human height, a negative control (b), and 14 immune-mediated diseases. Liability-scale *R*^2^ is shown for PRSs derived using ten significance thresholds (p_T_), represented by coloured bars. Significance estimates provided are for the least stringent p_T_<1 threshold, which included all SNPs. For complete results across representative p_T_, see **S3 Table**. Variance in schizophrenia case-status explained is approximately 0.8% for bipolar disorder PRSs using previously published data [11]. Variance in schizophrenia case-status explained by human height risk scores is approximately 0.06%, while immune-mediated disease PRSs explained <0.06% of variance. Disease abbreviations as defined in **S5 Table.**

**References**

1.	Chang CC, Chow CC, Tellier LC, Vattikuti S, Purcell SM, Lee JJ. Second-generation PLINK: rising to the challenge of larger and richer datasets. Gigascience. 2015;4:7.

2.	Bulik-Sullivan B, Finucane HK, Anttila V, Gusev A, Day FR, Loh PR, et al. An atlas of genetic correlations across human diseases and traits. Nat Genet. 2015;47(11):1236-41.

3.	Bulik-Sullivan BK, Loh PR, Finucane HK, Ripke S, Yang J, Patterson N, et al. LD Score regression distinguishes confounding from polygenicity in genome-wide association studies. Nat Genet. 2015;47(3):291-5.

4.	Power RA, Steinberg S, Bjornsdottir G, Rietveld CA, Abdellaoui A, Nivard MM, et al. Polygenic risk scores for schizophrenia and bipolar disorder predict creativity. Nat Neurosci. 2015;18(7):953-5.

5.	Stringer S, Kahn RS, de Witte LD, Ophoff RA, Derks EM. Genetic liability for schizophrenia predicts risk of immune disorders. Schizophr Res. 2014;159:347-52.

6.	Schizophrenia Working Group of the Psychiatric Genomics Consortium. Biological insights from 108 schizophrenia-associated genetic loci. Nature. 2014;511(7510):421-7.

7.	R Core Team. R: A language and environment for statistical computing. R Foundation for Statistical Computing. 2012;Vienna, Austria. ISBN 3-900051-07-0, URL http://www.R-project.org/.

8.	Lee SH, Goddard ME, Wray NR, Visscher PM. A better coefficient of determination for genetic profile analysis. Genet Epidemiol. 2012;36(3):214-24.

9.	Nakazawa M. fmsb: Functions for medical statistics book with some demographic data. R package version 051. 2014http://CRAN.R-project.org/package=fmsb.

10.	Purcell SM, Wray NR, Stone JL, Visscher PM, O’Donovan MC, Sullivan PF, et al. Common polygenic variation contributes to risk of schizophrenia and bipolar disorder. Nature. 2009;460(7256):748-52.

11.	Cross-Disorder Group of the Psychiatric Genomics Consortium. Identification of risk loci with shared effects on five major psychiatric disorders: a genome-wide analysis. Lancet. 2013;381:1371-9.

12.	Tylee DS, Sun J, Hess JL, Tahir MA, Sharma E, Malik R, et al. Genetic correlations among psychiatric and immune-related phenotypes based on genome-wide association data. Am J Med Genet B Neuropsychiatr Genet. 2018;177(7):641-57.

13.	Dubois PC, Trynka G, Franke L, Hunt KA, Romanos J, Curtotti A, et al. Multiple common variants for celiac disease influencing immune gene expression. Nat Genet. 2010;42(4):295-302.

14.	Wellcome Trust Case Control Consortium. Genome-wide association study of 14,000 cases of seven common diseases and 3,000 shared controls. Nature. 2007;447(7145):661-78.

15.	Cross-Disorder Group of the Psychiatric Genomics Consortium. Genetic relationship between five psychiatric disorders estimated from genome-wide SNPs. Nat Genet. 2013;45(9):984-94.

16.	Franke A, McGovern DP, Barrett JC, Wang K, Radford-Smith GL, Ahmad T, et al. Genome-wide meta-analysis increases to 71 the number of confirmed Crohn’s disease susceptibility loci. Nat Genet. 2010;42(12):1118-25.

17.	Liu JZ, van Sommeren S, Huang H, Ng SC, Alberts R, Takahashi A, et al. Association analyses identify 38 susceptibility loci for inflammatory bowel disease and highlight shared genetic risk across populations. Nat Genet. 2015;47(9):979-86.

18.	Wang Q, Yang C, Gelernter J, Zhao H. Pervasive pleiotropy between psychiatric disorders and immune disorders revealed by integrative analysis of multiple GWAS. Hum Genet. 2015;134(11-12):1195-209.

19.	Pickrell JK, Berisa T, Liu JZ, Segurel L, Tung JY, Hinds DA. Detection and interpretation of shared genetic influences on 42 human traits. Nat Genet. 2016;48(7):709-17.

20.	Jostins L, Ripke S, Weersma RK, Duerr RH, McGovern DP, Hui KY, et al. Host-microbe interactions have shaped the genetic architecture of inflammatory bowel disease. Nature. 2012;491(7422):119-24.

21.	Duncan LE, Shen H, Ballon JS, Hardy KV, Noordsy DL, Levinson DF. Genetic correlation profile of schizophrenia mirrors epidemiological results and suggests link between polygenic and rare variant (22q11.2) cases of schizophrenia. Schizophr Bull. 2018;44(6):1350-61.

22.	Andreassen OA, Harbo HF, Wang Y, Thompson WK, Schork AJ, Mattingsdal M, et al. Genetic pleiotropy between multiple sclerosis and schizophrenia but not bipolar disorder: differential involvement of immune-related gene loci. Mol Psychiatry. 2015;20:207-14.

23.	Sawcer S, Hellenthal G, Pirinen M, Spencer CC, Patsopoulos NA, Moutsianas L, et al. Genetic risk and a primary role for cell-mediated immune mechanisms in multiple sclerosis. Nature. 2011;476(7359):214-9.

24.	Ripke S, Sanders AR, Kendler KS, Levinson DF, Sklar P, Holmans PA, et al. Genome-wide association study identifies five new schizophrenia loci. Nat Genet. 2011;43(10):969-76.

25.	Hafler DA, Compston A, Sawcer S, Lander ES, Daly MJ, De Jager PL, et al. Risk alleles for multiple sclerosis identified by a genomewide study. N Engl J Med. 2007;357(9):851-62.

26.	Cordell HJ, Han Y, Mells GF, Li Y, Hirschfield GM, Greene CS, et al. International genome-wide meta-analysis identifies new primary biliary cirrhosis risk loci and targetable pathogenic pathways. Nat Commun. 2015;6:8019.

27.	Feng BJ, Sun LD, Soltani-Arabshahi R, Bowcock AM, Nair RP, Stuart P, et al. Multiple loci within the major histocompatibility complex confer risk of psoriasis. PLoS Genet. 2009;5(8):e1000606.

28.	Yin X, Wineinger NE, Wang K, Yue W, Norgren N, Wang L, et al. Common susceptibility variants are shared between schizophrenia and psoriasis in the Han Chinese population. J Psychiatry Neurosci. 2016;41(6):413-21.

29.	Yue WH, Wang HF, Sun LD, Tang FL, Liu ZH, Zhang HX, et al. Genome-wide association study identifies a susceptibility locus for schizophrenia in Han Chinese at 11p11.2. Nat Genet. 2011;43(12):1228-31.

30.	Zhang XJ, Huang W, Yang S, Sun LD, Zhang FY, Zhu QX, et al. Psoriasis genome-wide association study identifies susceptibility variants within LCE gene cluster at 1q21. Nat Genet. 2009;41(2):205-10.

31.	Tsoi LC, Spain SL, Knight J, Ellinghaus E, Stuart PE, Capon F, et al. Identification of 15 new psoriasis susceptibility loci highlights the role of innate immunity. Nat Genet. 2012;44(12):1341-8.

32.	Lee SH, Byrne EM, Hultman CM, Kahler A, Vinkhuyzen AA, Ripke S, et al. New data and an old puzzle: the negative association between schizophrenia and rheumatoid arthritis. Int J Epidemiol. 2015;44(5):1706-21.

33.	Ripke S, O’Dushlaine C, Chambert K, Moran JL, Kahler AK, Akterin S, et al. Genome-wide association analysis identifies 13 new risk loci for schizophrenia. Nat Genet. 2013;45(10):1150-9.

34.	Stahl EA, Raychaudhuri S, Remmers EF, Xie G, Eyre S, Thomson BP, et al. Genome-wide association study meta-analysis identifies seven new rheumatoid arthritis risk loci. Nat Genet. 2010;42(6):508-14.

35.	Okada Y, Wu D, Trynka G, Raj T, Terao C, Ikari K, et al. Genetics of rheumatoid arthritis contributes to biology and drug discovery. Nature. 2014;506(7488):376-81.

36.	Euesden J, Breen G, Farmer A, McGuffin P, Lewis CM. The relationship between schizophrenia and rheumatoid arthritis revisited: genetic and epidemiological analyses. Am J Med Genet B Neuropsychiatr Genet. 2015;168B(2):81-8.

37.	Harley JB, Alarcon-Riquelme ME, Criswell LA, Jacob CO, Kimberly RP, Moser KL, et al. Genome-wide association scan in women with systemic lupus erythematosus identifies susceptibility variants in ITGAM, PXK, KIAA1542 and other loci. Nat Genet. 2008;40(2):204-10.

38.	Bentham J, Morris DL, Cunninghame Graham DS, Pinder CL, Tombleson P, Behrens TW, et al. Genetic association analyses implicate aberrant regulation of innate and adaptive immunity genes in the pathogenesis of systemic lupus erythematosus. Nat Genet. 2015;47(12):1457-64.

39.	Radstake TR, Gorlova O, Rueda B, Martin JE, Alizadeh BZ, Palomino-Morales R, et al. Genome-wide association study of systemic sclerosis identifies CD247 as a new susceptibility locus. Nat Genet. 2010;42(5):426-9.

40.	Barrett JC, Clayton DG, Concannon P, Akolkar B, Cooper JD, Erlich HA, et al. Genome-wide association study and meta-analysis find that over 40 loci affect risk of type 1 diabetes. Nat Genet. 2009;41(6):703-7.

41.	Bradfield JP, Qu HQ, Wang K, Zhang H, Sleiman PM, Kim CE, et al. A genome-wide meta-analysis of six type 1 diabetes cohorts identifies multiple associated loci. PLoS Genet. 2011;7(9):e1002293.

42.	Anderson CA, Boucher G, Lees CW, Franke A, D’Amato M, Taylor KD, et al. Meta-analysis identifies 29 additional ulcerative colitis risk loci, increasing the number of confirmed associations to 47. Nat Genet. 2011;43(3):246-52.

43.	Schork AJ, Thompson WK, Pham P, Torkamani A, Roddey JC, Sullivan PF, et al. All SNPs are not created equal: genome-wide association studies reveal a consistent pattern of enrichment among functionally annotated SNPs. PLoS Genet. 2013;9(4):e1003449.

44.	Chung D, Yang C, Li C, Gelernter J, Zhao H. GPA: a statistical approach to prioritizing GWAS results by integrating pleiotropy and annotation. PLoS Genet. 2014;10(11):e1004787.

45.	Wray NR, Goddard ME, Visscher PM. Prediction of individual genetic risk to disease from genome-wide association studies. Genome Res. 2007;17(10):1520-8.

46.	Lee SH, Yang J, Goddard ME, Visscher PM, Wray NR. Estimation of pleiotropy between complex diseases using single-nucleotide polymorphism-derived genomic relationships and restricted maximum likelihood. Bioinformatics. 2012;28(19):2540-2.

47.	West J, Logan RF, Hill PG, Lloyd A, Lewis S, Hubbard R, et al. Seroprevalence, correlates, and characteristics of undetected coeliac disease in England. Gut. 2003;52(7):960-5.

48.	Trynka G, Hunt KA, Bockett NA, Romanos J, Mistry V, Szperl A, et al. Dense genotyping identifies and localizes multiple common and rare variant association signals in celiac disease. Nat Genet. 2011;43(12):1193-201.

49.	Stahl EA, Wegmann D, Trynka G, Gutierrez-Achury J, Do R, Voight BF, et al. Bayesian inference analyses of the polygenic architecture of rheumatoid arthritis. Nat Genet. 2012;44(5):483-9.

50.	Chen GB, Lee SH, Brion MJ, Montgomery GW, Wray NR, Radford-Smith GL, et al. Estimation and partitioning of (co)heritability of inflammatory bowel disease from GWAS and immunochip data. Hum Mol Genet. 2014;23(17):4710-20.

51.	Thompson SD, Marion MC, Sudman M, Ryan M, Tsoras M, Howard TD, et al. Genome-wide association analysis of juvenile idiopathic arthritis identifies a new susceptibility locus at chromosomal region 3q13. Arthritis Rheum. 2012;64(8):2781-91.

52.	Hinks A, Cobb J, Marion MC, Prahalad S, Sudman M, Bowes J, et al. Dense genotyping of immune-related disease regions identifies 14 new susceptibility loci for juvenile idiopathic arthritis. Nat Genet. 2013;45(6):664-9.

53.	Rosati G. The prevalence of multiple sclerosis in the world: an update. Neurol Sci. 2001;22(2):117-39.

54.	Beecham AH, Patsopoulos NA, Xifara DK, Davis MF, Kemppinen A, Cotsapas C, et al. Analysis of immune-related loci identifies 48 new susceptibility variants for multiple sclerosis. Nat Genet. 2013;45(11):1353-60.

55.	Watson CT, Disanto G, Breden F, Giovannoni G, Ramagopalan SV. Estimating the proportion of variation in susceptibility to multiple sclerosis captured by common SNPs. Sci Rep. 2012;2:770.

56.	Faraco J, Lin L, Kornum BR, Kenny EE, Trynka G, Einen M, et al. ImmunoChip study implicates antigen presentation to T cells in narcolepsy. PLoS Genet. 2013;9(2):e1003270.

57.	Liu JZ, Almarri MA, Gaffney DJ, Mells GF, Jostins L, Cordell HJ, et al. Dense fine-mapping study identifies new susceptibility loci for primary biliary cirrhosis. Nat Genet. 2012;44(10):1137-41.

58.	Parisi R, Symmons DP, Griffiths CE, Ashcroft DM. Global epidemiology of psoriasis: a systematic review of incidence and prevalence. J Invest Dermatol. 2013;133(2):377-85.

59.	Strange A, Capon F, Spencer CC, Knight J, Weale ME, Allen MH, et al. A genome-wide association study identifies new psoriasis susceptibility loci and an interaction between HLA-C and ERAP1. Nat Genet. 2010;42(11):985-90.

60.	Yin X, Wineinger NE, Cheng H, Cui Y, Zhou F, Zuo X, et al. Common variants explain a large fraction of the variability in the liability to psoriasis in a Han Chinese population. BMC Genomics. 2014;15:87.

61.	Bossini-Castillo L, Lopez-Isac E, Martin J. Immunogenetics of systemic sclerosis: Defining heritability, functional variants and shared-autoimmunity pathways. J Autoimmun. 2015;64:53-65.

62.	Sivertsen B, Petrie KJ, Wilhelmsen-Langeland A, Hysing M. Mental health in adolescents with Type 1 diabetes: results from a large population-based study. BMC Endocr Disord. 2014;14:83.

63.	Onengut-Gumuscu S, Chen WM, Burren O, Cooper NJ, Quinlan AR, Mychaleckyj JC, et al. Fine mapping of type 1 diabetes susceptibility loci and evidence for colocalization of causal variants with lymphoid gene enhancers. Nat Genet. 2015;47(4):381-6.

64.	Howitz J, Brodthagen H, Schwartz M, Thomsen K. Prevalence of vitiligo. Epidemiological survey on the Isle of Bornholm, Denmark. Arch Dermatol. 1977;113(1):47-52.

65.	Jin Y, Andersen G, Yorgov D, Ferrara TM, Ben S, Brownson KM, et al. Genome-wide association studies of autoimmune vitiligo identify 23 new risk loci and highlight key pathways and regulatory variants. Nat Genet. 2016;48(11):1418-24.

66.	Park JH, Wacholder S, Gail MH, Peters U, Jacobs KB, Chanock SJ, et al. Estimation of effect size distribution from genome-wide association studies and implications for future discoveries. Nat Genet. 2010;42(7):570-5.

67.	Gusev A, Bhatia G, Zaitlen N, Vilhjalmsson BJ, Diogo D, Stahl EA, et al. Quantifying missing heritability at known GWAS loci. PLoS Genet. 2013;9(12):e1003993.

68.	Pruim RJ, Welch RP, Sanna S, Teslovich TM, Chines PS, Gliedt TP, et al. LocusZoom: regional visualization of genome-wide association scan results. Bioinformatics. 2010;26(18):2336-7.

69.	Yang J, Lee SH, Goddard ME, Visscher PM. GCTA: a tool for genome-wide complex trait analysis. Am J Hum Genet. 2011;88(1):76-82.

70.	1000 Genomes Project Consortium, Auton A, Brooks LD, Durbin RM, Garrison EP, Kang HM, et al. A global reference for human genetic variation. Nature. 2015;526(7571):68-74.

71.	Dudbridge F. Power and predictive accuracy of polygenic risk scores. PLoS Genet. 2013;9(3):e1003348.

**Schizophrenia Working Group of the Psychiatric Genomics Consortium**

Stephan Ripke^1,2^, Benjamin M. Neale^1,2,3,4^, Aiden Corvin^5^, James T. R. Walters^6^, Kai-How Farh^1^, Peter A. Holmans^6,7^, Phil Lee^1,2,4^, Brendan Bulik-Sullivan^1,2^, David A. Collier^8,9^, Hailiang Huang^1,3^, Tune H. Pers^3,10,11^, Ingrid Agartz^12,13,14^, Esben Agerbo^15,16,17^, Margot Albus^18^, Madeline Alexander^19^, Farooq Amin^20,21^, Silviu A. Bacanu^22^, Martin Begemann^23^, Richard A Belliveau Jr^2^, Judit Bene^24,25^, Sarah E. Bergen ^2,26^, Elizabeth Bevilacqua^2^, Tim B Bigdeli ^22^, Donald W. Black^27^, Richard Bruggeman^28^, Nancy G. Buccola^29^, Randy L. Buckner^30,31,32^, William Byerley^33^, Wiepke Cahn^34^, Guiqing Cai^35,36^, Murray J. Cairns^39,120,170^, Dominique Campion^37^, Rita M. Cantor^38^, Vaughan J. Carr^39,40^, Noa Carrera^6^, Stanley V. Catts^39,41^, Kimberly D. Chambert^2^, Raymond C. K. Chan^42^, Ronald Y. L. Chen^43^, Eric Y. H. Chen^43,44^, Wei Cheng^45^, Eric F. C. Cheung^46^, Siow Ann Chong^47^, C. Robert Cloninger^48^, David Cohen^49^, Nadine Cohen^50^, Paul Cormican^5^, Nick Craddock^6,7^, Benedicto Crespo-Facorro^210^, James J. Crowley^51^, David Curtis^52,53^, Michael Davidson^54^, Kenneth L. Davis^36^, Franziska Degenhardt^55,56^, Jurgen Del Favero^57^, Lynn E. DeLisi^128,129^ , Ditte Demontis^17,58,59^, Dimitris Dikeos^60^, Timothy Dinan^61^, Srdjan Djurovic^14,62^, Gary Donohoe^5,63^, Elodie Drapeau^36^, Jubao Duan^64,65^, Frank Dudbridge^66^, Naser Durmishi^67^, Peter Eichhammer^68^, Johan Eriksson^69,70,71^, Valentina Escott-Price^6^, Laurent Essioux^72^, Ayman H. Fanous^73,74,75,76^, Martilias S. Farrell^51^, Josef Frank^77^, Lude Franke^78^, Robert Freedman^79^, Nelson B. Freimer^80^, Marion Friedl^81^, Joseph I. Friedman^36^, Menachem Fromer^1,2,4,82^, Giulio Genovese^2^, Lyudmila Georgieva^6^, Elliot S. Gershon^209^, Ina Giegling^81,83^, Paola Giusti-Rodríguez^51^, Stephanie Godard^84^, Jacqueline I. Goldstein^1,3^, Vera Golimbet^85^, Srihari Gopal^86^, Jacob Gratten^87^, Lieuwe de Haan^88^, Christian Hammer^23^, Marian L. Hamshere^6^, Mark Hansen^89^, Thomas Hansen^17,90^, Vahram Haroutunian^36,91,92^, Annette M. Hartmann^81^, Frans A. Henskens^39,93,94^, Stefan Herms^55,56,95^, Joel N. Hirschhorn^3,11,96^, Per Hoffmann^55,56,95^, Andrea Hofman^55,56^, Mads V. Hollegaard^97^, David M. Hougaard^97^, Masashi Ikeda^98^, Inge Joa^99^, Antonio Julià^100^, René S. Kahn^34^, Luba Kalaydjieva^101,102^, Sena Karachanak-Yankova^103^, Juha Karjalainen^78^, David Kavanagh^6^, Matthew C. Keller^104^, Brian J. Kelly^120^, James L. Kennedy^105,106,107^, Andrey Khrunin^108^, Yunjung Kim^51^, Janis Klovins^109^, James A. Knowles^110^, Bettina Konte^81^, Vaidutis Kucinskas^111^, Zita Ausrele Kucinskiene^111^, Hana Kuzelova-Ptackova^112^, Anna K. Kähler^26^, Claudine Laurent^19,113^, Jimmy Lee Chee Keong^47,114^, S. Hong Lee^87^, Sophie E. Legge^6^, Bernard Lerer^115^, Miaoxin Li^43,44,116^ Tao Li^117^, Kung-Yee Liang^118^, Jeffrey Lieberman^119^, Svetlana Limborska^108^, Carmel M. Loughland^39,120^, Jan Lubinski^121^, Jouko Lönnqvist^122^, Milan Macek Jr^112^, Patrik K. E. Magnusson^26^, Brion S. Maher^123^, Wolfgang Maier^124^, Jacques Mallet^125^, Sara Marsal^100^, Manuel Mattheisen^17,58,59,126^, Morten Mattingsdal^14,127^, Robert W. McCarley^128,129^, Colm McDonald^130^, Andrew M. McIntosh^131,132^, Sandra Meier^77^, Carin J. Meijer^88^, Bela Melegh^24,25^, Ingrid Melle^14,133^, Raquelle I. Mesholam-Gately^128,134^, Andres Metspalu^135^, Patricia T. Michie^39,136^, Lili Milani^135^, Vihra Milanova^137^, Younes Mokrab^8^, Derek W. Morris^5,63^, Ole Mors^17,58,138^, Kieran C. Murphy^139^, Robin M. Murray^140^, Inez Myin-Germeys^141^, Bertram Müller-Myhsok^142,143,144^, Mari Nelis^135^, Igor Nenadic^145^, Deborah A. Nertney^146^, Gerald Nestadt^147^, Kristin K. Nicodemus^148^, Liene Nikitina-Zake^109^, Laura Nisenbaum^149^, Annelie Nordin^150^, Eadbhard O’Callaghan^151^, Colm O’Dushlaine^2^, F. Anthony O’Neill^152^, Sang-Yun Oh^153^, Ann Olincy^79^, Line Olsen^17,90^, Jim Van Os^141,154^, Psychosis Endophenotypes International Consortium^155^, Christos Pantelis^39,156^, George N. Papadimitriou^60^, Sergi Papiol^23^, Elena Parkhomenko^36^, Michele T. Pato^110^, Tiina Paunio^157,158^, Milica Pejovic-Milovancevic^159^, Diana O. Perkins^160^, Olli Pietiläinen^158,161^, Jonathan Pimm^53^, Andrew J. Pocklington^6^, John Powell^140^, Alkes Price^3^,^162^, Ann E. Pulver^147^, Shaun M. Purcell^82^, Digby Quested^163^, Henrik B. Rasmussen^17,90^, Abraham Reichenberg^36^, Mark A. Reimers^164^, Alexander L. Richards^6^, Joshua L. Roffman^30,32^, Panos Roussos^82,165^, Douglas M. Ruderfer^6,82^, Veikko Salomaa^71^, Alan R. Sanders^64,65^, Ulrich Schall^39,120^, Christian R. Schubert^166^, Thomas G. Schulze^77,167^, Sibylle G. Schwab^168^, Edward M. Scolnick^2^, Rodney J. Scott^39,169,170^, Larry J. Seidman^128,134^, Jianxin Shi^171^, Engilbert Sigurdsson^172^, Teimuraz Silagadze^173^, Jeremy M. Silverman^36,174^, Kang Sim^47^, Petr Slominsky^108^, Jordan W. Smoller^2,4^, Hon-Cheong So^43^, Chris C. A. Spencer^175^, Eli A. Stahl^3,82^, Hreinn Stefansson^176^, Stacy Steinberg^176^, Elisabeth Stogmann^177^, Richard E. Straub^178^, Eric Strengman^179,34^, Jana Strohmaier^77^, T. Scott Stroup^119^, Mythily Subramaniam^47^, Jaana Suvisaari^122^, Dragan M. Svrakic^48^, Jin P. Szatkiewicz^51^, Erik Söderman^12^, Srinivas Thirumalai^180^, Draga Toncheva^103^, Paul A. Tooney^39,120,170^ , Sarah Tosato^181^, Juha Veijola^182,183^, John Waddington^184^, Dermot Walsh^185^, Dai Wang^86^, Qiang Wang^117^, Bradley T. Webb^22^, Mark Weiser^54^, Dieter B. Wildenauer^186^, Nigel M. Williams^6^, Stephanie Williams^51^, Stephanie H. Witt^77^, Aaron R. Wolen^164^, Emily H. M. Wong^43^, Brandon K. Wormley^22^, Jing Qin Wu^39,170^, Hualin Simon Xi^187^, Clement C. Zai^105,106^, Xuebin Zheng^188^, Fritz Zimprich^177^, Naomi R. Wray^87^, Kari Stefansson^176^, Peter M. Visscher^87^, Wellcome Trust Case-Control Consortium 2^189^, Rolf Adolfsson^150^, Ole A. Andreassen^14,133^, Douglas H. R. Blackwood^132^, Elvira Bramon^190^, Joseph D. Buxbaum^35,36,91,191^, Anders D. Børglum^17,58,59,138^, Sven Cichon^55,56,95,192^, Ariel Darvasi^193^, Enrico Domenici^194^, Hannelore Ehrenreich^23^, Tõnu Esko^3,11,96,135^, Pablo V. Gejman^64,65^, Michael Gill^5^, Hugh Gurling^53^, Christina M. Hultman^26^, Nakao Iwata^98^, Assen V. Jablensky^39,102,186,195^, Erik G. Jönsson^12,14^, Kenneth S. Kendler^196^, George Kirov^6^, Jo Knight^105,106,107^, Todd Lencz^197,198,199^, Douglas F. Levinson^19^, Qingqin S. Li^86^, Jianjun Liu^188,200^, Anil K. Malhotra^197,198,199^, Steven A. McCarroll^2,96^, Andrew McQuillin^53^, Jennifer L. Moran^2^, Preben B. Mortensen^15,16,17^, Bryan J. Mowry^87,201^, Markus M. Nöthen^55,56^, Roel A. Ophoff^38,80,34^, Michael J. Owen^6,7^, Aarno Palotie^2,4,161^, Carlos N. Pato^110^, Tracey L. Petryshen^2,128,202^, Danielle Posthuma^203,204,205^, Marcella Rietschel^77^, Brien P. Riley^196^, Dan Rujescu^81,83^, Pak C. Sham^43,44,116^ Pamela Sklar^82,91,165^, David St Clair^206^, Daniel R. Weinberger^178,207^, Jens R. Wendland^166^, Thomas Werge^17,90,208^, Mark J. Daly^1,2,3^, Patrick F. Sullivan^26,51,160^ & Michael C. O’Donovan^6,7^

^1^Analytic and Translational Genetics Unit, Massachusetts General Hospital, Boston, Massachusetts 02114, USA.

^2^Stanley Center for Psychiatric Research, Broad Institute of MIT and Harvard, Cambridge, Massachusetts 02142, USA.

^3^Medical and Population Genetics Program, Broad Institute of MIT and Harvard, Cambridge, Massachusetts 02142, USA.

^4^Psychiatric and Neurodevelopmental Genetics Unit, Massachusetts General Hospital, Boston, Massachusetts 02114, USA.

^5^Neuropsychiatric Genetics Research Group, Department of Psychiatry, Trinity College Dublin, Dublin 8, Ireland.

^6^MRC Centre for Neuropsychiatric Genetics and Genomics, Institute of Psychological Medicine and Clinical Neurosciences, School of Medicine, Cardiff University, Cardiff, CF24 4HQ, UK.

^7^National Centre for Mental Health, Cardiff University, Cardiff, CF24 4HQ, UK.

^8^Eli Lilly and Company Limited, Erl Wood Manor, Sunninghill Road, Windlesham, Surrey, GU20 6PH, UK. ^9^Social, Genetic and Developmental Psychiatry Centre, Institute of Psychiatry, King’s College London, London, SE5 8AF, UK.

^10^Center for Biological Sequence Analysis, Department of Systems Biology, Technical University of Denmark, DK-2800, Denmark.

^11^Division of Endocrinology and Center for Basic and Translational Obesity Research, Boston Children’s Hospital, Boston, Massachusetts, 02115USA.

^12^Department of Clinical Neuroscience, Psychiatry Section, Karolinska Institutet, SE-17176 Stockholm, Sweden. ^13^Department of Psychiatry, Diakonhjemmet Hospital, 0319 Oslo, Norway.

^14^NORMENT, KG Jebsen Centre for Psychosis Research, Institute of Clinical Medicine, University of Oslo, 0424 Oslo, Norway.

^15^Centre for Integrative Register-based Research, CIRRAU, Aarhus University, DK-8210 Aarhus, Denmark.

^16^National Centre for Register-based Research, Aarhus University, DK-8210 Aarhus, Denmark.

^17^The Lundbeck Foundation Initiative for Integrative Psychiatric Research, iPSYCH, Denmark.

^18^State Mental Hospital, 85540 Haar, Germany.

^19^Department of Psychiatry and Behavioral Sciences, Stanford University, Stanford, California 94305, USA.

^20^Department of Psychiatry and Behavioral Sciences, Atlanta Veterans Affairs Medical Center, Atlanta, Georgia 30033, USA.

^21^Department of Psychiatry and Behavioral Sciences, Emory University, Atlanta Georgia 30322, USA.

^22^Virginia Institute for Psychiatric and Behavioral Genetics, Department of Psychiatry, Virginia Commonwealth University, Richmond, Virginia 23298, USA.

^23^Clinical Neuroscience, Max Planck Institute of Experimental Medicine, Göttingen 37075, Germany.

^24^Department of Medical Genetics, University of Pécs, Pécs H-7624, Hungary.

^25^Szentagothai Research Center, University of Pécs, Pécs H-7624, Hungary.

^26^Department of Medical Epidemiology and Biostatistics, Karolinska Institutet, Stockholm SE-17177, Sweden.

^27^Department of Psychiatry, University of Iowa Carver College of Medicine, Iowa City, Iowa 52242, USA.

^28^University Medical Center Groningen, Department of Psychiatry, University of Groningen NL-9700 RB, The Netherlands.

^29^School of Nursing, Louisiana State University Health Sciences Center, New Orleans, Louisiana 70112, USA.

^30^Athinoula A. Martinos Center, Massachusetts General Hospital, Boston, Massachusetts 02129, USA.

^31^Center for Brain Science, Harvard University, Cambridge, Massachusetts, 02138 USA.

^32^Department of Psychiatry, Massachusetts General Hospital, Boston, Massachusetts, 02114 USA.

^33^Department of Psychiatry, University of California at San Francisco, San Francisco, California, 94143 USA.

^34^University Medical Center Utrecht, Department of Psychiatry, Rudolf Magnus Institute of Neuroscience, 3584 Utrecht, The Netherlands.

^35^Department of Human Genetics, Icahn School of Medicine at Mount Sinai, New York, New York 10029 USA.

^36^Department of Psychiatry, Icahn School of Medicine at Mount Sinai, New York, New York 10029 USA.

^37^Centre Hospitalier du Rouvray and INSERM U1079 Faculty of Medicine, 76301 Rouen, France.

^38^Department of Human Genetics, David Geffen School of Medicine, University of California, Los Angeles, California 90095, USA.

^39^Schizophrenia Research Institute, Sydney NSW 2010, Australia.

^40^School of Psychiatry, University of New South Wales, Sydney NSW 2031, Australia.

^41^Royal Brisbane and Women’s Hospital, University of Queensland, Brisbane, St Lucia QLD 4072, Australia.

^42^Institute of Psychology, Chinese Academy of Science, Beijing 100101, China.

^43^Department of Psychiatry, Li Ka Shing Faculty of Medicine, The University of Hong Kong, Hong Kong, China.

^44^State Key Laboratory for Brain and Cognitive Sciences, Li Ka Shing Faculty of Medicine, The University of Hong Kong, Hong Kong, China.

^45^Department of Computer Science, University of North Carolina, Chapel Hill, North Carolina 27514, USA.

^46^Castle Peak Hospital, Hong Kong, China.

^47^Institute of Mental Health, Singapore 539747, Singapore.

^48^Department of Psychiatry, Washington University, St. Louis, Missouri 63110, USA.

^49^Department of Child and Adolescent Psychiatry, Assistance Publique Hopitaux de Paris, Pierre and Marie Curie Faculty of Medicine and Institute for Intelligent Systems and Robotics, Paris, 75013, France.

^50^ Blue Note Biosciences, Princeton, New Jersey 08540, USA

^51^Department of Genetics, University of North Carolina, Chapel Hill, North Carolina 27599-7264, USA.

^52^Department of Psychological Medicine, Queen Mary University of London, London E1 1BB, UK.

^53^Molecular Psychiatry Laboratory, Division of Psychiatry, University College London, London WC1E 6JJ, UK.

^54^Sheba Medical Center, Tel Hashomer 52621, Israel.

^55^Department of Genomics, Life and Brain Center, D-53127 Bonn, Germany.

^56^Institute of Human Genetics, University of Bonn, D-53127 Bonn, Germany.

^57^Applied Molecular Genomics Unit, VIB Department of Molecular Genetics, University of Antwerp, B-2610 Antwerp, Belgium.

^58^Centre for Integrative Sequencing, iSEQ, Aarhus University, DK-8000 Aarhus C, Denmark.

^59^Department of Biomedicine, Aarhus University, DK-8000 Aarhus C, Denmark.

^60^First Department of Psychiatry, University of Athens Medical School, Athens 11528, Greece.

^61^Department of Psychiatry, University College Cork, Co. Cork, Ireland.

^62^Department of Medical Genetics, Oslo University Hospital, 0424 Oslo, Norway.

^63^Cognitive Genetics and Therapy Group, School of Psychology and Discipline of Biochemistry, National University of Ireland Galway, Co. Galway, Ireland.

^64^Department of Psychiatry and Behavioral Neuroscience, University of Chicago, Chicago, Illinois 60637, USA.

^65^Department of Psychiatry and Behavioral Sciences, NorthShore University HealthSystem, Evanston, Illinois 60201, USA.

^66^Department of Non-Communicable Disease Epidemiology, London School of Hygiene and Tropical Medicine, London WC1E 7HT, UK.

^67^Department of Child and Adolescent Psychiatry, University Clinic of Psychiatry, Skopje 1000, Republic of Macedonia.

^68^Department of Psychiatry, University of Regensburg, 93053 Regensburg, Germany.

^69^Department of General Practice, Helsinki University Central Hospital, University of Helsinki P.O. Box 20, Tukholmankatu 8 B, FI-00014, Helsinki, Finland

^70^Folkhälsan Research Center, Helsinki, Finland, Biomedicum Helsinki 1, Haartmaninkatu 8, FI-00290, Helsinki, Finland.

^71^National Institute for Health and Welfare, P.O. BOX 30, FI-00271 Helsinki, Finland.

^72^Translational Technologies and Bioinformatics, Pharma Research and Early Development, F. Hoffman-La Roche, CH-4070 Basel, Switzerland.

^73^Department of Psychiatry, Georgetown University School of Medicine, Washington DC 20057, USA.

^74^Department of Psychiatry, Keck School of Medicine of the University of Southern California, Los Angeles, California 90033, USA.

^75^Department of Psychiatry, Virginia Commonwealth University School of Medicine, Richmond, Virginia 23298, USA.

^76^Mental Health Service Line, Washington VA Medical Center, Washington DC 20422, USA.

^77^Department of Genetic Epidemiology in Psychiatry, Central Institute of Mental Health, Medical Faculty Mannheim, University of Heidelberg, Heidelberg , D-68159 Mannheim, Germany.

^78^Department of Genetics, University of Groningen, University Medical Centre Groningen, 9700 RB Groningen, The Netherlands.

^79^Department of Psychiatry, University of Colorado Denver, Aurora, Colorado 80045, USA.

^80^Center for Neurobehavioral Genetics, Semel Institute for Neuroscience and Human Behavior, University of California, Los Angeles, California 90095, USA.

^81^Department of Psychiatry, University of Halle, 06112 Halle, Germany.

^82^Division of Psychiatric Genomics, Department of Psychiatry, Icahn School of Medicine at Mount Sinai, New York, New York 10029, USA.

^83^Department of Psychiatry, University of Munich, 80336, Munich, Germany.

^84^Departments of Psychiatry and Human and Molecular Genetics, INSERM, Institut de Myologie, Hôpital de la Pitiè-Salpêtrière, Paris, 75013, France.

^85^Mental Health Research Centre, Russian Academy of Medical Sciences, 115522 Moscow, Russia.

^86^Neuroscience Therapeutic Area, Janssen Research and Development, Raritan, New Jersey 08869, USA.

^87^Queensland Brain Institute, The University of Queensland, Brisbane, Queensland, QLD 4072, Australia.

^88^Academic Medical Centre University of Amsterdam, Department of Psychiatry, 1105 AZ Amsterdam, The Netherlands.

^89^Illumina, La Jolla, California, California 92122, USA.

^90^Institute of Biological Psychiatry, Mental Health Centre Sct. Hans, Mental Health Services Copenhagen, DK-4000, Denmark.

^91^Friedman Brain Institute, Icahn School of Medicine at Mount Sinai, New York, New York 10029, USA.

^92^J. J. Peters VA Medical Center, Bronx, New York, New York 10468, USA.

^93^Priority Research Centre for Health Behaviour, University of Newcastle, Newcastle NSW 2308, Australia.

^94^School of Electrical Engineering and Computer Science, University of Newcastle, Newcastle NSW 2308, Australia.

^95^Division of Medical Genetics, Department of Biomedicine, University of Basel, Basel, CH-4058, Switzerland.

^96^Department of Genetics, Harvard Medical School, Boston, Massachusetts 02115, USA.

^97^Section of Neonatal Screening and Hormones, Department of Clinical Biochemistry, Immunology and Genetics, Statens Serum Institut, Copenhagen, DK-2300, Denmark.

^98^Department of Psychiatry, Fujita Health University School of Medicine, Toyoake, Aichi, 470-1192, Japan.

^99^Regional Centre for Clinical Research in Psychosis, Department of Psychiatry, Stavanger University Hospital, 4011 Stavanger, Norway.

^100^Rheumatology Research Group, Vall d'Hebron Research Institute, Barcelona, 08035, Spain.

^101^Centre for Medical Research, The University of Western Australia, Perth, WA 6009, Australia.

^102^The Perkins Institute for Medical Research, The University of Western Australia, Perth, WA 6009, Australia.

^103^Department of Medical Genetics, Medical University, Sofia1431, Bulgaria.

^104^Department of Psychology, University of Colorado Boulder, Boulder, Colorado 80309, USA.

^105^Campbell Family Mental Health Research Institute, Centre for Addiction and Mental Health, Toronto, Ontario, M5T 1R8, Canada.

^106^Department of Psychiatry, University of Toronto, Toronto, Ontario, M5T 1R8, Canada.

^107^Institute of Medical Science, University of Toronto, Toronto, Ontario, M5S 1A8, Canada.

^108^Institute of Molecular Genetics, Russian Academy of Sciences, Moscow123182, Russia.

^109^Latvian Biomedical Research and Study Centre, Riga, LV-1067, Latvia.

^110^Department of Psychiatry and Zilkha Neurogenetics Institute, Keck School of Medicine at University of Southern California, Los Angeles, California 90089, USA.

^111^Faculty of Medicine, Vilnius University, LT-01513 Vilnius, Lithuania.

^112^ Department of Biology and Medical Genetics, 2nd Faculty of Medicine and University Hospital Motol, 150 06 Prague, Czech Republic.

^113^ Department of Child and Adolescent Psychiatry, Pierre and Marie Curie Faculty of Medicine, Paris 75013, France.

^114^Duke-NUS Graduate Medical School, Singapore 169857, Singapore.

^115^Department of Psychiatry, Hadassah-Hebrew University Medical Center, Jerusalem 91120, Israel.

^116^Centre for Genomic Sciences, The University of Hong Kong, Hong Kong, China.

^117^Mental Health Centre and Psychiatric Laboratory, West China Hospital, Sichuan University, Chengdu, 610041, Sichuan, China.

^118^Department of Biostatistics, Johns Hopkins University Bloomberg School of Public Health, Baltimore, Maryland 21205, USA.

^119^Department of Psychiatry, Columbia University, New York, New York 10032, USA.

^120^Priority Centre for Translational Neuroscience and Mental Health, University of Newcastle, Newcastle NSW 2300, Australia.

^121^Department of Genetics and Pathology, International Hereditary Cancer Center, Pomeranian Medical University in Szczecin, 70-453 Szczecin, Poland.

^122^Department of Mental Health and Substance Abuse Services; National Institute for Health and Welfare, P.O. BOX 30, FI-00271 Helsinki, Finland

^123^Department of Mental Health, Bloomberg School of Public Health, Johns Hopkins University, Baltimore, Maryland 21205, USA.

^124^Department of Psychiatry, University of Bonn, D-53127 Bonn, Germany.

^125^Centre National de la Recherche Scientifique, Laboratoire de Génétique Moléculaire de la Neurotransmission et des Processus Neurodégénératifs, Hôpital de la Pitié Salpêtrière, 75013, Paris, France.

^126^Department of Genomics Mathematics, University of Bonn, D-53127 Bonn, Germany.

^127^Research Unit, Sørlandet Hospital, 4604 Kristiansand, Norway.

^128^Department of Psychiatry, Harvard Medical School, Boston, Massachusetts 02115, USA.

^129^VA Boston Health Care System, Brockton, Massachusetts 02301, USA.

^130^Department of Psychiatry, National University of Ireland Galway, Co. Galway, Ireland.

^131^Centre for Cognitive Ageing and Cognitive Epidemiology, University of Edinburgh, Edinburgh EH16 4SB, UK.

^132^Division of Psychiatry, University of Edinburgh, Edinburgh EH16 4SB, UK.

^133^Division of Mental Health and Addiction, Oslo University Hospital, 0424 Oslo, Norway.

^134^Massachusetts Mental Health Center Public Psychiatry Division of the Beth Israel Deaconess Medical Center, Boston, Massachusetts 02114, USA.

^135^Estonian Genome Center, University of Tartu, Tartu 50090, Estonia.

^136^School of Psychology, University of Newcastle, Newcastle NSW 2308, Australia.

^137^First Psychiatric Clinic, Medical University, Sofia 1431, Bulgaria.

^138^Department P, Aarhus University Hospital, DK-8240 Risskov, Denmark.

^139^Department of Psychiatry, Royal College of Surgeons in Ireland, Dublin 2, Ireland.

^140^King’s College London, London SE5 8AF, UK.

^141^Maastricht University Medical Centre, South Limburg Mental Health Research and Teaching Network, EURON, 6229 HX Maastricht, The Netherlands.

^142^Institute of Translational Medicine, University of Liverpool, Liverpool L69 3BX, UK.

^143^Max Planck Institute of Psychiatry, 80336 Munich, Germany.

^144^Munich Cluster for Systems Neurology (SyNergy), 80336 Munich, Germany.

^145^Department of Psychiatry and Psychotherapy, Jena University Hospital, 07743 Jena, Germany.

^146^Department of Psychiatry, Queensland Brain Institute and Queensland Centre for Mental Health Research, University of Queensland, Brisbane, Queensland, St Lucia QLD 4072, Australia.

^147^Department of Psychiatry and Behavioral Sciences, Johns Hopkins University School of Medicine, Baltimore, Maryland 21205, USA.

^148^Department of Psychiatry, Trinity College Dublin, Dublin 2, Ireland.

^149^Eli Lilly and Company, Lilly Corporate Center, Indianapolis, 46285 Indiana, USA.

^150^Department of Clinical Sciences, Psychiatry, Umeå University, SE-901 87 Umeå, Sweden.

^151^DETECT Early Intervention Service for Psychosis, Blackrock, Co. Dublin, Ireland.

^152^Centre for Public Health, Institute of Clinical Sciences, Queen’s University Belfast, Belfast BT12 6AB, UK.

^153^Lawrence Berkeley National Laboratory, University of California at Berkeley, Berkeley, California 94720, USA.

^154^Institute of Psychiatry, King’s College London, London SE5 8AF, UK.

^155^A list of authors and affiliations appear in the Supplementary Information.

^156^Melbourne Neuropsychiatry Centre, University of Melbourne & Melbourne Health, Melbourne, Vic 3053, Australia.

^157^Department of Psychiatry, University of Helsinki, P.O. Box 590, FI-00029 HUS, Helsinki, Finland.

^158^Public Health Genomics Unit, National Institute for Health and Welfare, P.O. BOX 30, FI-00271 Helsinki, Finland.

^159^Medical Faculty, University of Belgrade, 11000 Belgrade, Serbia.

^160^Department of Psychiatry, University of North Carolina, Chapel Hill, North Carolina 27599-7160, USA.

^161^Institute for Molecular Medicine Finland, FIMM, University of Helsinki, P.O. Box 20

FI-00014, Helsinki, Finland.

^162^Department of Epidemiology, Harvard School of Public Health, Boston, Massachusetts 02115, USA.

^163^Department of Psychiatry, University of Oxford, Oxford, OX3 7JX, UK.

^164^Virginia Institute for Psychiatric and Behavioral Genetics, Virginia Commonwealth University, Richmond, Virginia 23298, USA.

^165^Institute for Multiscale Biology, Icahn School of Medicine at Mount Sinai, New York, New York 10029, USA.

^166^PharmaTherapeutics Clinical Research, Pfizer Worldwide Research and Development, Cambridge, Massachusetts 02139, USA.

^167^Department of Psychiatry and Psychotherapy, University of Gottingen, 37073 Göttingen, Germany.

^168^Psychiatry and Psychotherapy Clinic, University of Erlangen, 91054 Erlangen, Germany.

^169^Hunter New England Health Service, Newcastle NSW 2308, Australia.

^170^School of Biomedical Sciences and Pharmacy, University of Newcastle, Callaghan NSW 2308, Australia.

^171^Division of Cancer Epidemiology and Genetics, National Cancer Institute, Bethesda, Maryland 20892, USA.

^172^University of Iceland, Landspitali, National University Hospital, 101 Reykjavik, Iceland.

^173^Department of Psychiatry and Drug Addiction, Tbilisi State Medical University (TSMU), **N33, 0177** Tbilisi, Georgia.

^174^Research and Development, Bronx Veterans Affairs Medical Center, New York, New York 10468, USA.

^175^Wellcome Trust Centre for Human Genetics, Oxford, OX3 7BN, UK.

^176^deCODE Genetics, 101 Reykjavik, Iceland.

^177^Department of Clinical Neurology, Medical University of Vienna, 1090 Wien, Austria.

^178^Lieber Institute for Brain Development, Baltimore, Maryland 21205, USA.

^179^Department of Medical Genetics, University Medical Centre Utrecht, Universiteitsweg 100, 3584 CG, Utrecht, The Netherlands.

^180^Berkshire Healthcare NHS Foundation Trust, Bracknell RG12 1BQ, UK.

^181^Section of Psychiatry, University of Verona, 37134 Verona, Italy.

^182^Department of Psychiatry, University of Oulu, P.O. BOX 5000, 90014, Finland

^183^University Hospital of Oulu, P.O.BOX 20, 90029 OYS, Finland.

^184^Molecular and Cellular Therapeutics, Royal College of Surgeons in Ireland, Dublin 2, Ireland.

^185^Health Research Board, Dublin 2, Ireland.

^186^School of Psychiatry and Clinical Neurosciences, The University of Western Australia, Perth WA6009, Australia.

^187^Computational Sciences CoE, Pfizer Worldwide Research and Development, Cambridge, Massachusetts 02139, USA.

^188^Human Genetics, Genome Institute of Singapore, A*STAR, Singapore 138672, Singapore.

^189^A list of authors and affiliations appear in the Supplementary Information.

^190^University College London, London WC1E 6BT, UK.

^191^Department of Neuroscience, Icahn School of Medicine at Mount Sinai, New York, New York 10029, USA.

^192^Institute of Neuroscience and Medicine (INM-1), Research Center Juelich, 52428 Juelich, Germany.

^193^Department of Genetics, The Hebrew University of Jerusalem, 91905 Jerusalem, Israel.

^194^Neuroscience Discovery and Translational Area, Pharma Research and Early Development, F. Hoffman-La Roche, CH-4070 Basel, Switzerland.

^195^Centre for Clinical Research in Neuropsychiatry, School of Psychiatry and Clinical Neurosciences, The University of Western Australia, Medical Research Foundation Building, Perth WA 6000, Australia.

^196^Virginia Institute for Psychiatric and Behavioral Genetics, Departments of Psychiatry and Human and Molecular Genetics, Virginia Commonwealth University, Richmond, Virginia 23298, USA.

^197^The Feinstein Institute for Medical Research, Manhasset, New York, 11030 USA.

^198^The Hofstra NS-LIJ School of Medicine, Hempstead, New York, 11549 USA.

^199^The Zucker Hillside Hospital, Glen Oaks, New York,11004 USA.

^200^Saw Swee Hock School of Public Health, National University of Singapore, Singapore 117597, Singapore.

^201^Queensland Centre for Mental Health Research, University of Queensland, Brisbane 4076, Queensland, Australia.

^202^Center for Human Genetic Research and Department of Psychiatry, Massachusetts General Hospital, Boston, Massachusetts 02114, USA.

^203^Department of Child and Adolescent Psychiatry, Erasmus University Medical Centre, Rotterdam 3000, The Netherlands.

^204^Department of Complex Trait Genetics, Neuroscience Campus Amsterdam, VU University Medical Center Amsterdam, Amsterdam 1081, The Netherlands.

^205^Department of Functional Genomics, Center for Neurogenomics and Cognitive Research, Neuroscience Campus Amsterdam, VU University, Amsterdam 1081, The Netherlands.

^206^University of Aberdeen, Institute of Medical Sciences, Aberdeen, AB25 2ZD, UK.

^207^Departments of Psychiatry, Neurology, Neuroscience and Institute of Genetic Medicine, Johns Hopkins School of Medicine, Baltimore, Maryland 21205, USA.

^208^Department of Clinical Medicine, University of Copenhagen, Copenhagen 2200, Denmark.

^209^Departments of Psychiatry and Human Genetics, University of Chicago, Chicago, Illinois 60637, USA.

^210^University Hospital Marqués de Valdecilla, Instituto de Formación e Investigación Marqués de Valdecilla, University of Cantabria, E‐39008 Santander, Spain.
